# Peripheral phoneme encoding and discrimination in aging and hearing impairment

**DOI:** 10.64898/2026.01.27.702044

**Authors:** Marjoleen Wouters, Etienne Gaudrain, Konrad Dapper, Jakob Schirmer, Deniz Başkent, Lukas Rüttiger, Marlies Knipper, Sarah Verhulst

## Abstract

**Highlights:**

- Speech EEG shows age- and OHC-related vulnerability of high-CF envelope coding.
- Speech EEG shows low-CF envelope encoding stays intact with age.
- Fine-structure contrast discrimination worsens with OHC loss in quiet.
- Fine-structure contrast discrimination worsens with age in contralateral noise.
- Phoneme discrimination involving high-frequency contrasts remains robust with age.
- Peripheral coding strength is not directly reflected at behavioral level.

Speech perception difficulties in noise are common among older adults and individuals with hearing impairment, but also occur among younger adults whose audiometric thresholds are within the clinically-normal range. We examined how aging, cochlear synaptopathy (CS), and outer hair cell (OHC) damage affect speech encoding and phoneme discrimination. Envelope-following responses (EFRs) to rectangular amplitude-modulated (RAM) tones and speech-like phoneme pairs were recorded in quiet using EEG, and behavioral discrimination was assessed in quiet, and with ipsilateral and contralateral noise. Stimuli were designed to target temporal envelope (TENV) or temporal fine structure (TFS) encoding. Results showed that RAM-EFR amplitudes decreased gradually with increasing age, consistent with emerging CS, while magnitudes of high-frequency TENV-based EFRs in quiet were most reduced in older hearing-impaired subjects with combined CS and OHC damage. In contrast, EFRs targeting low-frequency TENV encoding in quiet remained preserved. Behaviorally, discrimination of predominantly TFS-based phoneme contrasts worsened with OHC loss and age in quiet and contralateral noise, respectively, whereas discrimination of predominantly TENV-based contrasts showed no significant age effect. Considering that high-frequency contrasts are discriminated via place-based spectral cues, low-frequency contrasts rely on TFS, and the EFR reflects primarily TENV, this framework explains why EFRs decline for high-frequency cues without perceptual loss, while EFRs remain stable for low-frequency cues even as TFS-based discrimination deteriorates. These findings highlight the need to investigate how neural coding deficits relate to perception. Combining electrophysiological and behavioral measures might provide a framework for detecting subclinical auditory deficits to earlier diagnose age-related and hidden hearing loss.

## 1. Introduction

Speech understanding in noisy environments is a common complaint among individuals with hearing difficulties, and is often associated with age-related or acquired sensorineural hearing loss (SNHL) (Furman et al., 2013; Kujawa and Liberman, 2009; Lobarinas et al., 2017). A well-established hallmark of SNHL is outer hair cell (OHC) damage, which reduces cochlear gain and elevates hearing thresholds on the pure-tone audiogram, the standard diagnostic tool in clinical audiology (Moore, 2007). However, mounting evidence suggests that impaired speech-in-noise perception can also occur in the absence of elevated audiometric thresholds, pointing to additional mechanisms beyond OHC damage (Plack et al., 2014; Liberman, 2017; Oxenham, 2016; Brungart et al., 2022).

One such other aspect of SNHL is cochlear synaptopathy (CS), a pathology that has often been associated with so-called “hidden hearing loss”. CS is characterized by degeneration of the synapses between inner hair cells (IHCs) and auditory nerve (AN) fibers, resulting in a reduced neural representation of sound without elevation of audiometric pure-tone thresholds at conventional audiometric frequencies (“normal” hearing is defined by the World Health Organization as pure-tone threshold average of the frequencies 0.5, 1, 2 and 4 kHz (PTA4) ≤ 25 dB HL) (Kujawa and Liberman, 2009; Sergeyenko et al., 2013; Liberman et al., 2015; Möhrle et al., 2016, 2019; Humes, 2018). CS is commonly associated with aging, noise exposure, and ototoxic drug use, and may precede OHC degeneration (Furman et al., 2013; Bramhall et al., 2017). Given these features, CS is often considered a subclinical hearing disorder, referring to auditory deficits that arise before changes become evident on standard audiometry, despite clinically “normal” auditory thresholds within conventional frequency ranges (e.g. below 4 kHz; (Liberman et al., 2016; Skoe and Tufts, 2018)). In CS, low-spontaneous-rate (LSR) AN fibers are typically affected first, particularly in noise-exposure and aging models, although the pattern of vulnerability can vary across species (Furman et al., 2013; Schmiedt et al., 1996). These fibers have high thresholds and are crucial for encoding the properties of sounds in noisy environments, such as amplitude modulation. Because they are more vulnerable to noise exposure and aging, their synapses are often lost before those of high-SR fibers, even when conventional audiometric thresholds remain “normal” (Kujawa and Liberman, 2009, 2015; Furman et al., 2013; Schmiedt et al., 1996). Consequently, individuals with clinically normal audiograms may still experience substantial difficulties understanding speech in noise, which can negatively impact cognition, psychosocial well-being, and educational outcomes, yet frequently remain undiagnosed and untreated (Bharadwaj et al., 2015; Plack et al., 2014; Bakay et al., 2018).

Electrophysiological measures have proven valuable in identifying neural markers of CS. In particular, the envelope-following response (EFR) evoked by rectangular amplitude-modulated (RAM) pure tones has been proposed as a sensitive correlate of synaptic integrity (Vasilkov et al., 2021; Keshishzadeh et al., 2021; Vasilkov and Verhulst, 2019). The RAM-EFR has been shown to be largely insensitive to other aspects of SNHL such as OHC damage, since the sharply rising envelope of the RAM stimulus causes a large, synchronized neural response almost instantly, which minimizes the time for the cochlear amplifier to function (Keshishzadeh et al., 2020, 2021; Vasilkov et al., 2021; Van Der Biest et al., 2023).

Additionally, speech-evoked auditory evoked potentials (AEPs) allow for assessment of temporal coding over different frequency ranges: low-frequency temporal fine structure (TFS) coding below the assumed phase-locking limit (PLL, around 1.5 kHz ^1^), and high-frequency temporal envelope (TENV) coding above the PLL (Lorenzi et al., 2006; Verschooten et al., 2019). By comparing neural and behavioral responses to phoneme contrasts presented in quiet and different noise configurations, it is possible to probe how CS, OHC damage, and aging differentially affect speech encoding.

In the present study, we extended our previous work (Schirmer et al., 2024; Dapper et al., 2025) by investigating the relationship between speech encoding and phoneme discrimination across the lifespan and in the presence of hearing impairment. We focused on disentangling the effects of aging, CS, and OHC damage on (i) RAM-EFR magnitude, (ii) speech-evoked EEG responses to phoneme contrasts above and below the PLL, and (iii) behavioral phoneme discrimination in quiet and in noise. Our subject groups included young normal-hearing (yNH), middle-aged normal-hearing (mNH), older normal-hearing (oNH), and older hearing-impaired (oHI) adults.

We hypothesized that: (i) CS and aging would be reflected in reduced RAM-EFR and speech EEG responses, even for NH groups, (ii) phoneme discrimination, particularly in ipsilateral noise, would be impaired in individuals with CS, with consonant confusions being most prominent, and (iii) OHC damage would further exacerbate deficits in oHI individuals. We expected that the yNH group would serve as a baseline control, with robust AEPs, intact speech encoding and phoneme discrimination in quiet and noise, and no CS or central gain changes (increased neural responsiveness in central auditory pathways due to SNHL in the cochlea) (Möhrle et al., 2016; Sergeyenko et al., 2013; Parthasarathy and Kujawa, 2018). For the mNH group, we expected to see subtle age-related CS emerge (Möhrle et al., 2016; Sergeyenko et al., 2013; Parthasarathy and Kujawa, 2018), with possible central gain compensation (Schirmer et al., 2024). Speech coding was expected to worsen more in noise than in quiet, likely due to reduced sampling by AN fibers, because when fewer AN fibers carry detailed information, the neural representation of speech becomes less precise, especially in challenging listening situations (Lopez-Poveda and Barrios, 2013). For the oNH group, we expected even stronger age-related effects on AEPs and speech encoding, particularly in noise. Although central gain mechanisms may partially compensate for the reduced peripheral encoding associated with CS, they are unlikely to restore normal performance in speech encoding (Sergeyenko et al., 2013; Auerbach et al., 2014; Möhrle et al., 2016). Finally, for the oHI group, we expected combined effects of OHC damage, age-related CS, and central gain changes, with strongly degraded speech AEPs, and worse speech discrimination in quiet and noise.

By combining EEG markers of peripheral speech encoding with phoneme discrimination performance across carefully stratified age and hearing-status groups, we aimed to differentiate the contributions of aging, CS, and OHC damage to speech perception deficits. Identifying reliable neural markers for subclinical hearing disorders will be essential for advancing early diagnosis and targeted interventions.

## 2. Materials and Methods

### 2.1. Ethics statement

“The study was conducted in the Department of Otolaryngology of the University of Tübingen, and approved by the ethics committee of Tübingen University (Faculty of Medicine; ethical approval number 392/2021BO2). Written, informed consent was given by all participants. All methods followed the Declaration of Helsinki by the World Medical Association (WMA) for human research ethics.” (Schirmer et al., 2024)

### 2.2. Subject selection

In this study, we selected a subset of 49 out of the 112 subjects who were tested in the study of Schirmer et al. (2024), based on their age and audiogram. Within each group, the age range was limited in order to neglect the effect of age. Audiometric hearing thresholds were assessed at 11 frequencies ranging from 0.125 to 10 kHz, as well as at 4 frequencies for the extended high-frequency range of 11.2 to 16 kHz (Schirmer et al., 2024). The pure-tone threshold average across the frequencies 0.5, 1, 2 and 4 kHz (PTA4) was calculated for each subject. Subjects were divided into five groups: yNH1 (18-24 years old, n = 10, 5 females), yNH2 (25-31 years old, n = 10, 5 females), mNH (32-48 years old, n = 9, 7 females), oNH (49-65 years old, n = 10, 7 females), oHI (49-65 years old, n = 10, 8 females, with worse PTA4 than oNH). Figure 1 shows the audiometric thresholds for all five groups, as well as their age distribution.

**Figure 1:**
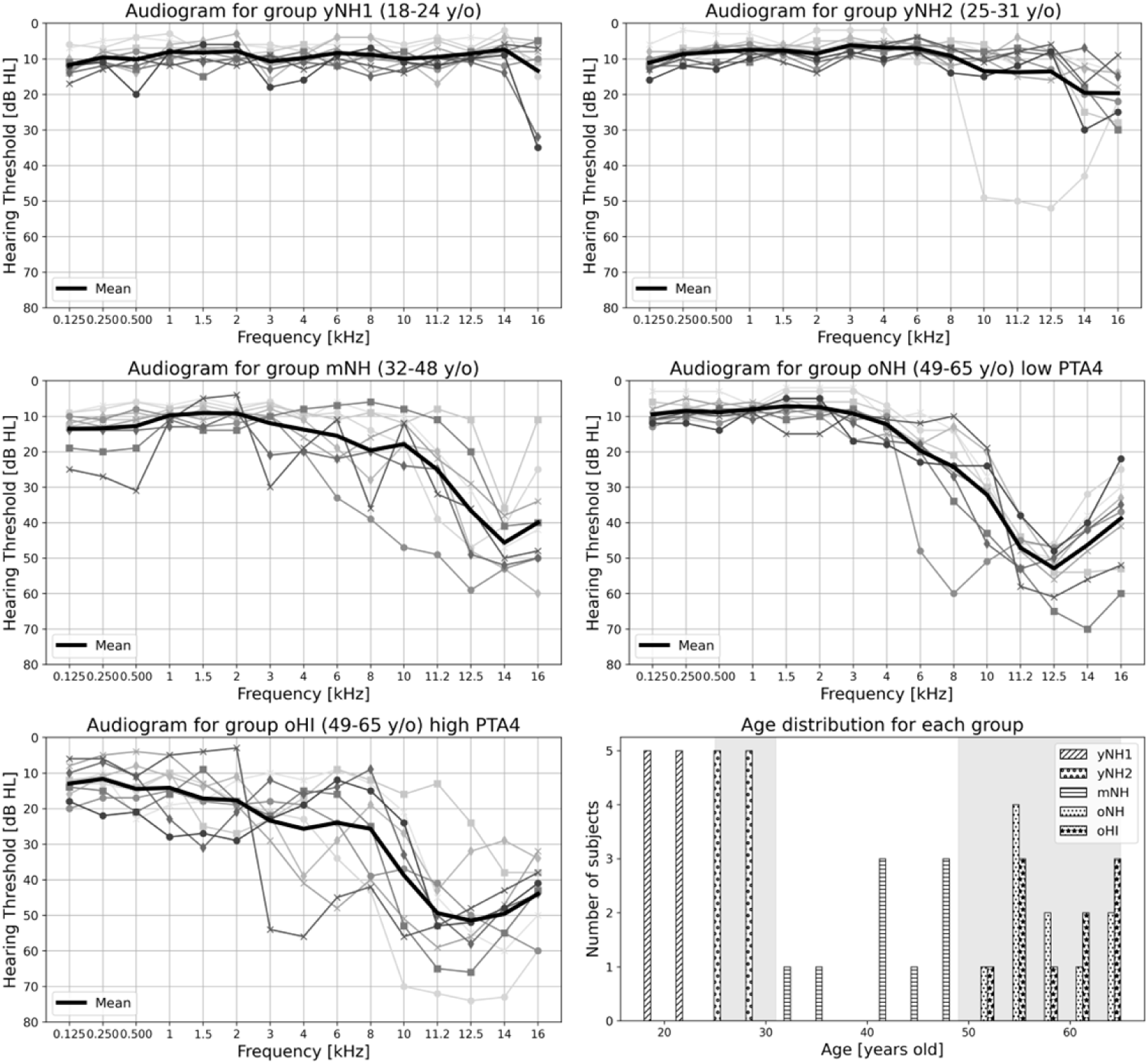
Audiograms and age histogram for each group. In the age histogram, grey background bands were added to visually separate the groups.

Subjects in the first four groups were selected from their age group based on the lowest PTA4 values, all below 20 dB HL and hence considered “normal” (Humes, 2018; Von Gablenz and Holube, 2017). Subjects in the group oHI were selected to have relatively higher PTA4 values within their age group (≥ 17 dB HL). Although this cutoff reflects only mild elevations in auditory thresholds, it was chosen to ensure a sufficient sample size, while still representing the upper end of the PTA4 distribution. Despite the modest PTA4 elevation, the oHI group exhibited substantially greater degrees of cochlear gain loss relative to the oNH group according to their audiogram. The PTA4 values for each group are shown in Figure 2a using boxplots with overlaid individual data points as solid black circles, and outliers displayed as open circles. The Shapiro-Wilk test, implemented in Python (v.3.9.13), showed deviation from normality for the oHI group (*W* = .6202, *p* = .0001). The Kruskal-Wallis H test (*H* = 28.370, *p* ≤ .001), followed by Dunn’s post hoc comparisons with Holm correction, showed a significantly higher PTA4 value for group oHI than for the other groups (yNH1: *p* < .001, yNH2: *p* < .001 and oNH: *p* < .01). The number of subjects in each group was limited because of the requirement for good PTA4 values (below 20 dB HL for the NH groups), exclusion criteria (Schirmer et al., 2024), and the availability of complete recordings of their speech EEG.

**Figure 2:**
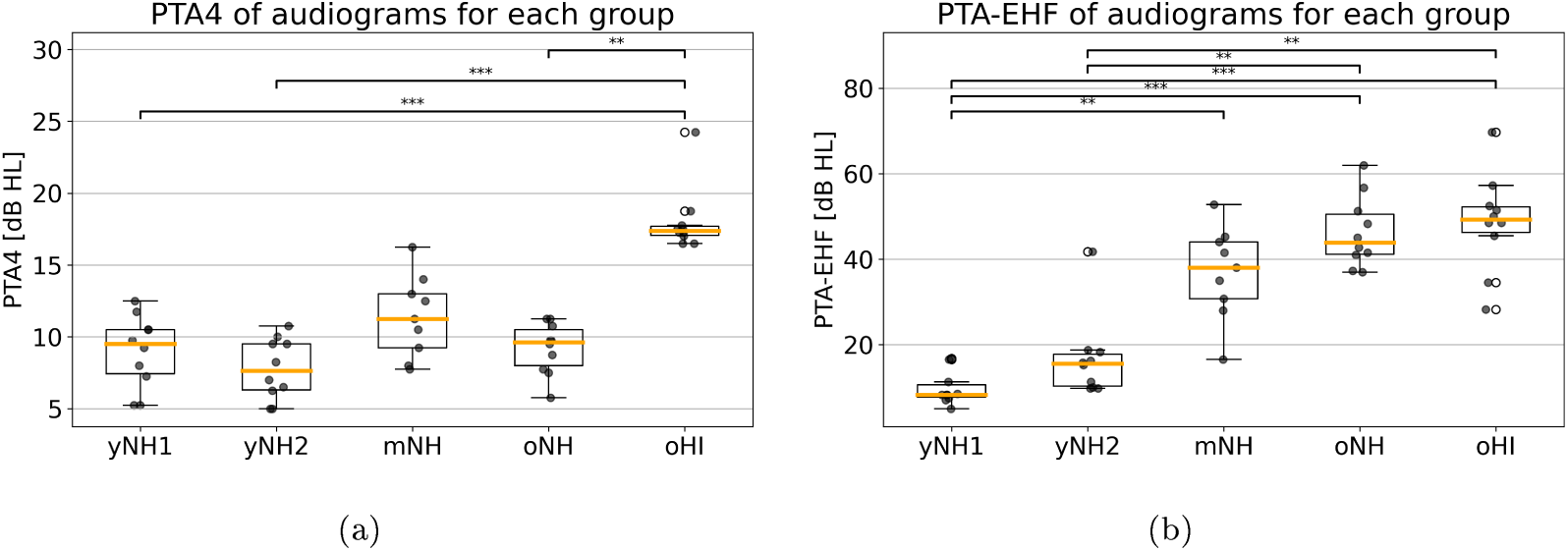
Group comparisons of PTA4 (a) and PTA-EHF (b) (*p* < .001:***; *p* < .01:**; *p <* .05:*).

For all five groups, we calculated the pure-tone threshold average for the extended high-frequencies (PTA-EHF), namely 11.2, 12.5, 14 and 16 kHz, which are known to be particularly vulnerable to neural degeneration (Bramhall et al., 2017; Lobarinas et al., 2013; Liberman et al., 2016), while OHC dysfunction may also be especially pronounced in this frequency range (Wang et al., 2021; Wu et al., 2019). Figure 2b shows the PTA-EHF values for each group. To investigate differences in PTA-EHF values across the groups, a Shapiro-Wilk test was first conducted to assess the normality of the data within each group. The results indicated that the PTA-EHF values for two groups (yNH1: *W* = .8161, *p* = .0228; yNH2: *W* = .6960, *p* = .0008) significantly deviated from normality, while the values for the remaining groups did not *(p* > .05). Given this violation of normality, a Kruskal-Wallis H test was employed. The test revealed a significant difference in PTA-EHF thresholds across groups (*H* = 34.977, *p* < .0001), suggesting that at least one group differed from the others. Pairwise comparisons were conducted using Dunn’s test with Holm correction to control for multiple comparisons. The results are shown in Figure 2b using boxplots with overlaid individual data points and annotated significance bars. PTA-EHF values for group yNHl were significantly lower than those for groups mNH *(p* < .01), oNH *(p* < .001) and oHI *(p <* .001). PTA-EHF values for group yNH2 were significantly lower than those for groups oNH (*p <* .01) and oHI *(p <* .01). This is consistent with previous studies showing that hearing thresholds in the EHF range worsen with age and frequency, with measurable declines emerging already during adolescence and becoming more pronounced from approximately 31-40 years onward (Wang et al., 2021; Green et al., 1987). There was no significant difference in PTA-EHF values between the mNH, oNH and oHI groups.

### 2.3. Phoneme stimuli

We generated four pairs of phoneme stimuli using the analysis-resynthesis framework of the WORLD vocoder (Morise et al., 2016), following the same approach as described in Schirmer et al. (2024) and Dapper et al. (2025), to be used in a phoneme-discrimination task and to investigate peripheral speech encoding of those phonemes through speech EEG (Gaudrain et al., 2025). All stimuli were synthesized using a male voice with a fundamental frequency (*F*_0_) of 116 Hz to match the OLSA sentences (German Oldenburger sentence test; Wagener et al. (1999)), and a sampling frequency of 44.1 kHz. Stimuli were constructed such that some pairs differed only in spectral regions below the assumed PLL of 1.5 kHz, thereby allowing discrimination predominantly through TFS cues: /du/-/bu/ (as in “<u>doe</u>” versus “<u>boe</u>te” in Dutch) and /o/-/u/ (as in “b<u>oo</u>t” versus “<u>oe</u>ster” in Dutch). The other pairs differed only above the assumed PLL, restricting discrimination predominantly to TENV cues: /di/-/bi/ (as in “die” versus “bier” in Dutch) and /i/-/y/ (as in “<u>ie</u>ts” versus “<u>u</u>niversiteit” in Dutch) (Weiss and Rose, 1988; Gaudrain et al., 2023, 2025). It is worth noting that the upper frequency limit of neural phase-locking in humans remains uncertain, with estimates ranging from approximately 1.5 to 10 kHz (Verschooten et al., 2019). Accordingly, we do not treat 1.5 kHz as a strict physiological cutoff but rather as a pragmatic reference for the present analyses, based on the binaural PLL of TFS coding (Verschooten et al., 2019) and on cue availability in speech (Ardoint and Lorenzi, 2010). Previous work suggests that TFS coding is most robust at low frequencies and progressively degrades with increasing frequency, with complete loss likely occurring only at substantially higher frequencies (Verschooten et al., 2019; Klug et al., 2023). Thus, frequencies below 1.5 kHz are expected to support reliable TFS cues, whereas above this range their contribution may be reduced but not absent, with TENV cues becoming increasingly dominant. Moreover, the 1-2 kHz region encompasses a peak in the speech-importance function (Yoho et al., 2018), implying that frequencies immediately below and above 1.5 kHz both contribute meaningfully to speech perception.

The steady vowel segments were standardized across syllable pairs, such that the steady state of the single vowel phonemes corresponded to the vowel part of the corresponding consonant-vowel syllables from 100 ms onward (e.g. the steady vowel /i/ was matched to the steady state /i/ segment embedded in /di/ and /bi/). Stimuli were synthesized with 30-ms onset/offset ramps and the total stimulus durations were 170 ms for the vowel pairs and 230 ms for the syllable pairs (Schirmer et al., 2024; Dapper et al., 2025; Gaudrain et al., 2025).

Finally, the stimuli were spectrally shaped to ensure comparable effective signal-to-noise ratios (SNRs) above and below the PLL relative to the OLSA speech-shaped noise (Gaudrain et al., 2025). The stimuli were then level-equalized such that the average level within each stimulus pair was identical across pairs and equal to 60 dB SPL (*L_eq_*), while preserving minor level differences within pairs to avoid altering formant structure. Calibration was performed using an IEC 60318-4 ear simulator (Schirmer et al., 2024; Dapper et al., 2025). Figure 3 shows the time-domain waveforms of all stimulus pairs. The consonants /d/ and /b/ occurred at about 50 - 60 ms, and after 100 ms there was no difference between the vowel segments of the syllable pairs (Gaudrain et al., 2025). Figure 3 also shows the frequency spectra and Figure 4 shows the short-time Fourier transform (STFT) of all stimuli. The STFT was calculated using a Hann window with a size of 500 samples, number of fast Fourier transform points (NFFT) = 500 samples and hop length = 120 samples, with a sampling frequency of FS = 44.1 kHz. In the STFT spectrograms, sound levels are displayed as short-time spectral magnitudes in dB SPL, referenced to the same calibration as the overall stimulus level; thus, peak values approach 60 dB SPL while lower values reflect the temporal and spectral distribution of energy within the signal. From Figures 3 and 4, it can be seen that the contrast of the /o/-/u/ pair was in the lower harmonics of *F*_0_ = 116 Hz. The envelope of /o/ was higher in amplitude than that of /u/, but the latter had a higher magnitude of carrier peaks within a lower envelope. For the /i/-/y/ pair, the contrast was between 2-4 kHz in the 2nd, 3rd and 4th formant frequencies (as shown in Table 1 in Dapper et al. (2025)). The contrast for the /di/-/bi/ pair was in the 2nd formant around 2.4 - 2.6 kHz, and for the /du/-/bu/ pair the contrast was in the first and second formants around 0.8 - 1.2 kHz, as shown in Figure 1B in Gaudrain et al. (2025).

**Figure 3:**
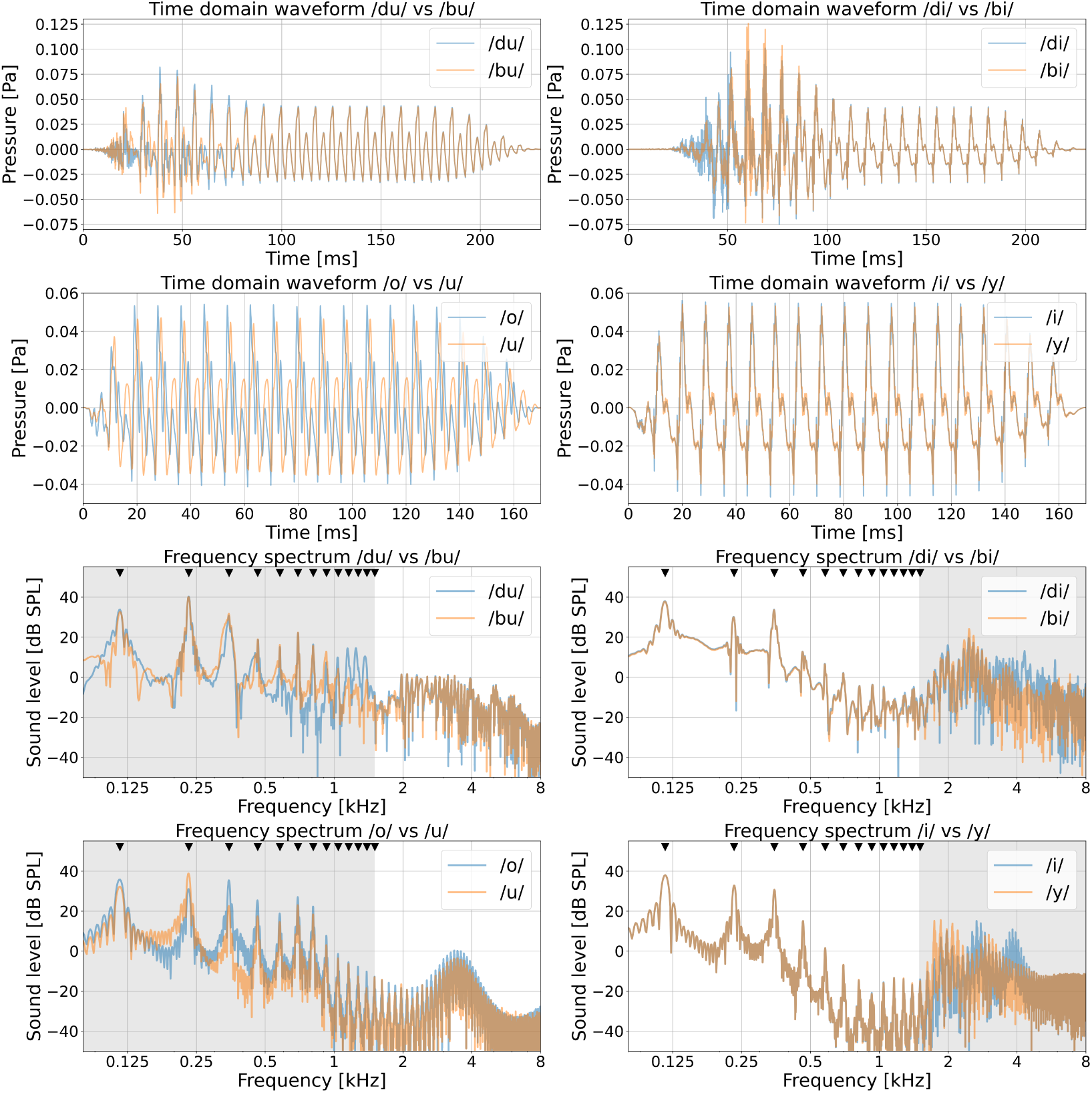
Time-domain waveforms and frequency spectra of the stimuli. The harmonics of *F*_0_ are marked by black triangle symbols, and the grey background area marks the frequency range in which there is a contrast within the stimulus pair (below or above the assumed PLL).

**Figure 4:**
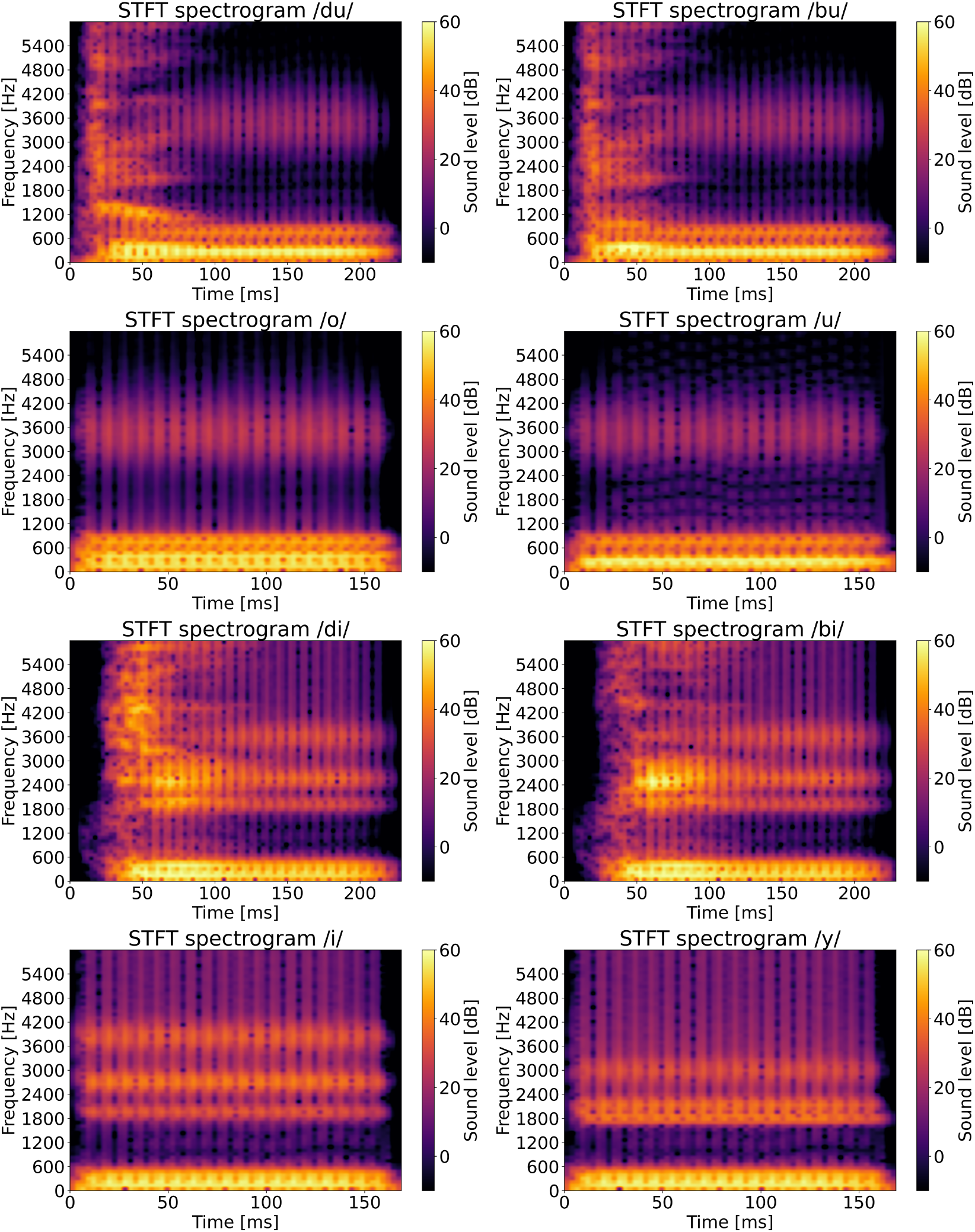
STFT spectrograms of the stimuli (FS = 44.1 kHz, window shape = Hann, window size = 500 samples, NFFT = 500 samples, hop length = 120 samples).

### 2.4. Speech EEG recording

For the speech EEG recording, the speech-like stimuli (/di/, /bi/, /du/, /bu/, /o/ and /y/) were presented binaurally in quiet at 60 dB SPL in an interleaved stimuli protocol, using the methodology and equipment set-up described by Dapper et al. (2025). Each stimulus was zero-padded to an epoch length of 250 ms, jittered in time by +/- 5 ms, and presented 3456 times, with alternating epochs of positive and negative acoustic waveform polarity (to cancel the effect of the cochlear microphonic) via insert earphones (ER1-14A) and mu-metal shielded ER2 transducers (Etymotic Research, Elk Grove Village, Illinois, USA). The subject lay on their back in a horizontal position inside an electrically and acoustically shielded booth (Dapper et al., 2025). Recordings were performed using passive stick-on electrodes (Neuroline 720, Ambu, Bad Nauheim, Germany) positioned on both mastoids, a reference electrode at the Fpz position below the hairline, and a ground electrode in between the eyebrows, with a sampling rate of FS = 50 kHz for each EEG channel (Dapper et al., 2025). Due to time constraints, the vowels /u/ and /i/ were not used during the speech EEG session, since their responses could be extracted from the recordings of the syllables /du/ and /di/, respectively (Dapper et al., 2025).

Although neural responses cannot be attributed exclusively to a single anatomical level, the speech-evoked EFRs measured in this study are interpreted as predominantly reflecting processing at the level of the auditory brainstem. When using the traditional clinical vertical electrode montage, EFRs elicited by modulation frequencies in the range of 80 - 100 Hz have been shown to be mainly generated by brainstem and midbrain sources (Herdman et al., 2002). Cortical contributions to the EFR mainly occur at lower modulation rates (Coffey et al., 2016, 2017; Bidelman, 2018; Herdman et al., 2002).

### 2.5. Pre- and post-processing of EEG data

The raw data of the EFRs were pre-processed using Python (v.3.9.13) and the MNE-Python package (v.0.23.4) (Gramfort et al., 2013, 2014). First, the Fpz channel recording was re-referenced to the average of the two mastoid electrodes. Second, the triggers of the EEG epochs were extracted to identify the stimulus to which the EFR corresponded. Then, artifacts were removed from the EEG data by applying a band-pass filter between 60 and 1500 Hz, and epochs of length 250 ms (−15 to 235 ms) were extracted based on the trigger events, including 1728 positive and 1728 negative polarity epochs per stimulus. Next, the epochs were baseline-corrected. Then, for each polarity the epochs with the highest peak-to-peak amplitudes, which might correspond to artifacts, were rejected, such that 1650 epochs with the lowest peak-to-peak amplitude remained for each polarity. To obtain the time-domain waveform of the EFR to each stimulus, the 3300 epochs were averaged.

#### Frequency domain EFR analysis

For the analysis of the EEG recordings in the frequency domain, we calculated the fast Fourier transform (FFT) for each of the epochs using NFFT = 50,000 bins, to get a frequency resolution of 1 Hz for the EFR spectrum (*F_res_ = FS/NFFT).* For /u/ and /i/ we cut off the first 60 ms of the EFR to /du/ and /di/ respectively. We cut off the last 60 ms of the EFR response to /o/ and /y/, to have an equal FFT length for the pairs /o/-/u/ and /i/-/y/. We used a bootstrapping procedure of random sampling with replacement, for 500 iterations, to obtain the EFR frequency spectrum of each stimulus with noise. To derive the noise floor, we performed a bootstrapping procedure for 500 iterations on the EFR spectra of each stimulus, using the phase flip method proposed by Zhu et al. (2013). For each bootstrap draw, randomly selected epochs from the recording channel were split into two equal groups, with one group phase-inverted (multiplied by −1). Averaging the combined epochs removes any consistently phase-locked neural signal, because responses with opposite phase cancel each other, while random noise (which is not phase-aligned) remains. Repeating this procedure across iterations yields null responses, which are EFRs that contain only noise and no real EFR signal, which were used to derive a channel-specific noise floor. Finally, we subtracted the noise floor spectrum from the full EFR spectrum, and averaged it over the 500 iterations, to acquire the clean noise floor-corrected EFR spectrum for each stimulus. To illustrate this, Figure 5 shows the EFR spectrum in response to the stimulus /bi/ for one subject from group yNHl.

**Figure 5:**
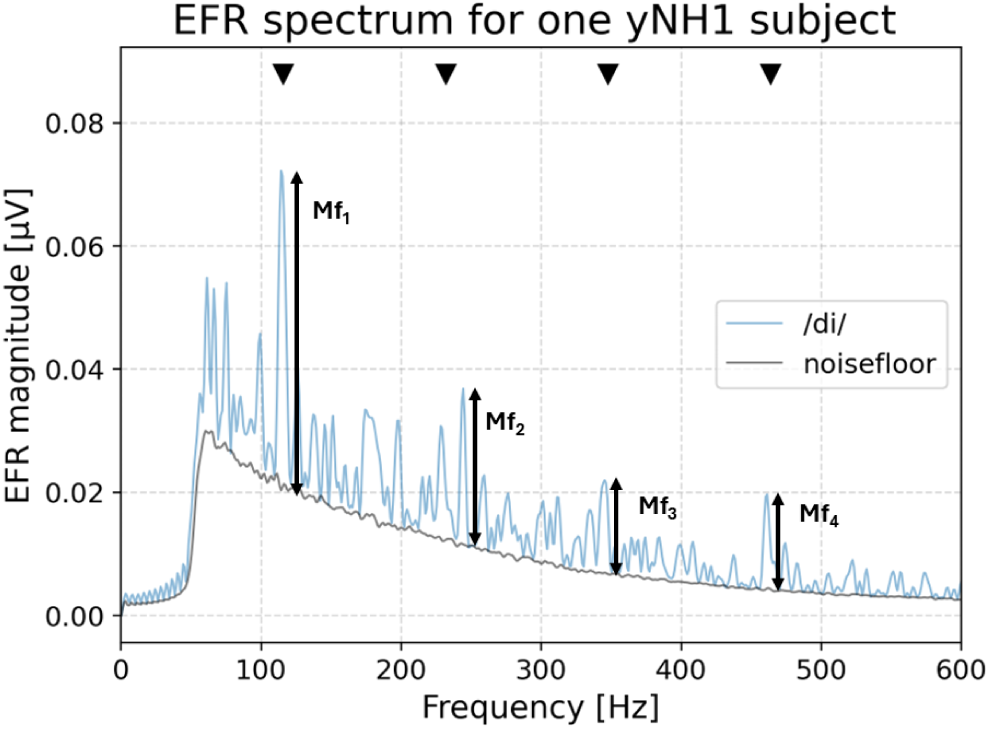
EFR frequency spectrum for the /bi/ stimulus for one subject from group yNHl. The harmonics of *F*_0_ are marked by black triangle symbols. *M_f1_-M_f4_* represent the EFR amplitudes of the four harmonics of *F*_0_.

From this noise floor-corrected EFR spectrum, we calculated for each subject the EFR peak value as the mean of the EFR amplitudes *M_fk_* of the four harmonics of the fundamental frequency *F*_0_ = 116 Hz, according to Equation 1.

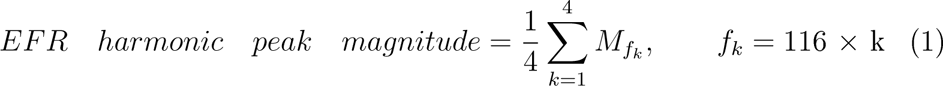

To evaluate the effects of stimulus type on EFR magnitude, we conducted both between-group and within-group statistical analyses in Python. For the between-group comparisons, EFR magnitudes were averaged for each stimulus and assessed for normality using the Shapiro-Wilk test. If the data for all groups met the assumption of normality, a one-way ANOVA was performed followed by post hoc pairwise comparisons using Tukey’s HSD test to identify significant differences. In cases where normality was violated, the non-parametric Kruskal-Wallis test was applied, and pair-wise comparisons were conducted using Dunn’s post hoc test with Holm correction. For within-group comparisons, data for paired stimuli were analyzed using either the paired t-test or the Wilcoxon signed-rank test, depending on the normality of the difference scores. Multiple comparisons were corrected using the Benjamini-Hochberg procedure to control the false discovery rate. All statistical outcomes were visualized using boxplots with individual data points and significance markers.

For each stimulus, we investigated the correlation between EFR magnitude and age using linear regression analysis. The EFR magnitude values were paired with the subject’s age. The slope (*β*) quantified the rate of magnitude change with age, the coefficient of determination (*R*^2^) assessed the proportion of variance explained by the model, and statistical significance was evaluated using the associated p-value. Pooled correlations were computed across all groups for each stimulus.

#### Time-domain EFR analysis

To assess the similarity in neural responses to different phoneme pairs, we computed normalized maximum cross-correlations between averaged EFR time-domain waveforms for each group. Higher correlation means that responses to the two stimuli are more similar. Before calculating the cross-correlation, we calculated the time shift of the stimulus time-domain waveform in response to /u/ from that for /du/ and /i/ from that for /di/, so as to match the peaks in the waveforms of /o/ and /y/, respectively, and applied the same shift to the EFR time-domain waveforms. For each phoneme pair, we segmented the EFR time-domain waveform into consonant and vowel time windows using a 50-kHz sampling rate. The consonant window spanned the initial 100 ms (Gaudrain et al., 2025), while vowel windows varied depending on the phoneme duration (for /du/-/bu/ and /di/-/bi/: 100 - 250 ms; for /o/-/u/ and /i/-/y/: 50 - 170 ms). The cross-correlation was calculated only within these time windows, to isolate phoneme-specific temporal dynamics. The similarity between the specific time windows of two stimuli in each pair was quantified for each subject using the maximum value of the normalized cross-correlation function. This analysis was performed separately for each group, using the available number of subjects per group, yielding correlation values for consonant and vowel segments that enabled comparison across conditions.

To assess group-level differences in neural similarity of the EFR time-domain waveform for each phoneme pair, we conducted separate statistical analyses for consonant and vowel segments based on cross-correlation values. For each segment type, data were grouped by phoneme pair and first tested for normality using the Shapiro-Wilk test. If all groups satisfied the assumption of normality, a one-way ANOVA was performed to evaluate overall group differences, followed by Tukey’s HSD post hoc tests for specific pairwise contrasts. When normality was violated, the non-parametric Kruskal-Wallis test was applied, with post hoc comparisons conducted using Dunn’s test with Holm correction. These procedures were applied independently to both consonant and vowel data.

### 2.6. EFR to a RAM pure tone stimulus

Using the same measurement setup as for the speech EEG recordings, we recorded the EFR to a RAM pure tone with a 4-kHz carrier frequency (RAM-4000), modulation frequency of 116 Hz and modulation depth of 100%, using 400 positive and 400 negative polarity epochs of length 500 ms (stimulus length 400 ms) at a sound level of 70 dB SPL rms and with a sampling rate of 50 kHz. The EFR to a RAM-4000 stimulus is a sensitive marker of CS, for which lower EFR magnitudes correspond to higher degrees of CS (Vasilkov et al., 2021; Keshishzadeh et al., 2021). Although the RAM stimulus is designed to have a sharply rising envelope, auditory filtering will inevitably smear this onset at the level of the basilar membrane. Due to the band-pass properties of cochlear filtering, a sudden change in stimulus amplitude cannot be represented instantaneously, but is temporally smeared such that the effective envelope rise spans several cycles of the carrier frequency (Verschooten et al., 2019; Evans, 1989). As a result, some transient engagement of the cochlear amplifier and associated gain changes may occur during this brief onset period. Nevertheless, compared to more slowly modulated stimuli, the rapid envelope modulation is expected to substantially reduce sustained OHC-related gain effects on the steady state RAM-EFR (Van Der Biest et al., 2023; Keshishzadeh et al., 2021; Vasilkov et al., 2021).

From the RAM-EFR recordings, the EFR peak magnitudes were calculated for each subject using the Matlab (v.2022b) toolbox EEGLAB (v.2021.1) The EEG signals were band-pass filtered between 60 and 1000 Hz, epoched, baseline-corrected, and the 200 epochs with the highest peak-to-peak values were rejected. The frequency spectra were calculated using Matlab’s FFT function for NFFT = 50,000 bins, to obtain a frequency resolution of 1 Hz. Then, a bootstrapping procedure using the phase flip method was executed using 400 draws to separate the noise floor from the clean EFR spectrum. The magnitude of the RAM-EFR of each of the 400 draws was calculated from the noise floor-corrected spectrum as the sum of the frequency response at the first four harmonics of the fundamental frequency *F*_0_ = 116 Hz, and then averaged to obtain one RAM-EFR magnitude value per subject.

To assess the significance of differences in RAM-EFR magnitude across subject groups, a non-parametric Kruskal-Wallis rank sum test was performed in Python. The Kruskal-Wallis test was selected due to a significant violation of the normality assumption required for parametric ANOVA, as indicated by the Shapiro-Wilk test (mNH: *W* = .8079, *p* = .0251). The correlation between RAM-EFR magnitude and age was assessed using linear regression analysis. RAM-EFR magnitudes were paired with each subject’s age, and a linear model was fitted to estimate the association between age and RAM-EFR magnitude.

### 2.7. Phoneme discrimination task

We employed the same phoneme discrimination paradigm as described by Schirmer et al. (2024), using a three-alternative forced-choice (3AFC) task with the four phoneme pairs described earlier. Subjects were presented with three sequentially delivered phonemes for each phoneme pair, two of which were identical and one different. The task was to identify the phoneme that differed from the other two by indicating it on the computer screen. To allow systematic adjustment of task difficulty, the phoneme stimuli were generated along a nine-step continuum ranging from one end of each phonemic contrast to the other (e.g. from /o/ to /u/) (Gaudrain et al., 2025). Task difficulty was quantified using these nine levels, which were evaluated in pilot experiments to identify stimulus pairs with predefined levels of behavioral discriminability. For the psychoacoustic session, two stimulus pairs per phoneme contrast were selected: one easy and one difficult, corresponding to a larger and smaller separation within the continuum, respectively. Such flexibility is particularly important in studies including HI subjects, where NH subjects may perform at “ceiling” while HI subjects may perform at “floor” level.

The discrimination task was performed in quiet, with ipsilateral noise, and with contralateral noise, to dissociate the effects of peripheral modulation masking from those of binaural and central auditory processing under different spatial configurations. Speech intelligibility in noise is known to be largely determined by modulation masking rather than purely energetic masking, even for background noise that is notionally steady (Stone et al., 2012; Stone and Moore, 2014; Stone et al., 2011). Ipsilateral noise was used to introduce modulation masking by directly interfering with speech TENV cues. In contrast, contralateral noise does not acoustically overlap with the target signal at the stimulated ear and therefore does not induce peripheral masking; instead, it introduces competing modulation patterns across ears, thereby primarily engaging binaural integration and central auditory processing mechanisms (Wendt et al., 2021).

To reduce learning effects, the order of the conditions was randomized, and a brief initial training phase was included with visual feedback, consisting of four phoneme discrimination trials for each of the four phoneme pairs. This training was intended to familiarize subjects with the task and response format rather than to induce learning. Nine repetitions were completed for each stimulus pair, yielding 54 trials in total. Stimuli were presented over ER2 transducers to the right ear. The noise was speech-shaped noise taken from the OLSA test, presented at 0 dB SNR. For each stimulus pair and noise condition, phoneme discrimination scores were expressed on a scale from 0 to 1. Scores below 0.33 indicate performance below chance level.

Between-group differences for each stimulus pair and noise condition and within-group differences across the noise conditions and stimulus pairs were examined using two sets of pairwise comparisons and generalized linear mixed-effects models (GLMMs) with a binomial link function in R (v.4.5.1, lme4 v.1.1.37). First, within each group, we assessed the effect of noise presence and location (quiet, ipsilateral, contralateral) across stimulus pairs by fitting GLMMs that included noise location and stimulus pair as fixed effects, and subject nested within difficulty level as a random effect to account for repeated measures. Estimated marginal means (emmeans v.1.11.2) were computed for each noise location within each stimulus pair, and pairwise contrasts were used to determine whether specific configurations of noise significantly affected the phoneme discrimination scores. Second, to examine between-group differences, we performed pairwise comparisons of response accuracy across groups for each combination of stimulus pair and noise location. For each subset, GLMMs were fitted with group as a fixed effect and subject nested within difficulty level as a random effect. Estimated marginal means and pairwise contrasts were used to identify significant differences in performance between groups under matched listening conditions. This dual modeling approach allowed us to disentangle the effects of noise configuration and group membership on phoneme discrimination performance.

## 3. Results

### 3.1. EFR in the frequency domain

The RAM-EFR peak values for each subject group are shown in Figure 6a, using boxplots with overlaid individual data points as filled black circles, and outliers displayed as open circles. The Kruskal-Wallis H test revealed a marginally significant difference in EFR magnitudes across the groups *(H =* 10.338, *p* = .0351), but Dunn’s post hoc comparisons with Holm correction did not show a significant difference between groups. This likely reflects a combination of substantial within-group variability and comparable magnitudes for the yNHl and yNH2 groups, and for the oNH and oHI groups. As seen in Figure 6a, the yNH1 group contained a clear outlier. This point was retained for non-parametric comparisons, but excluded from the linear regression due to the sensitivity of least-squares methods to outliers. Figure 6b shows a significant negative correlation of RAM-EFR magnitude with age (*β* = −0.0017, *R*^2^ = .161, *p* = .005), indicating that increasing age was associated with reduced RAM-EFR magnitude.

**Figure 6:**
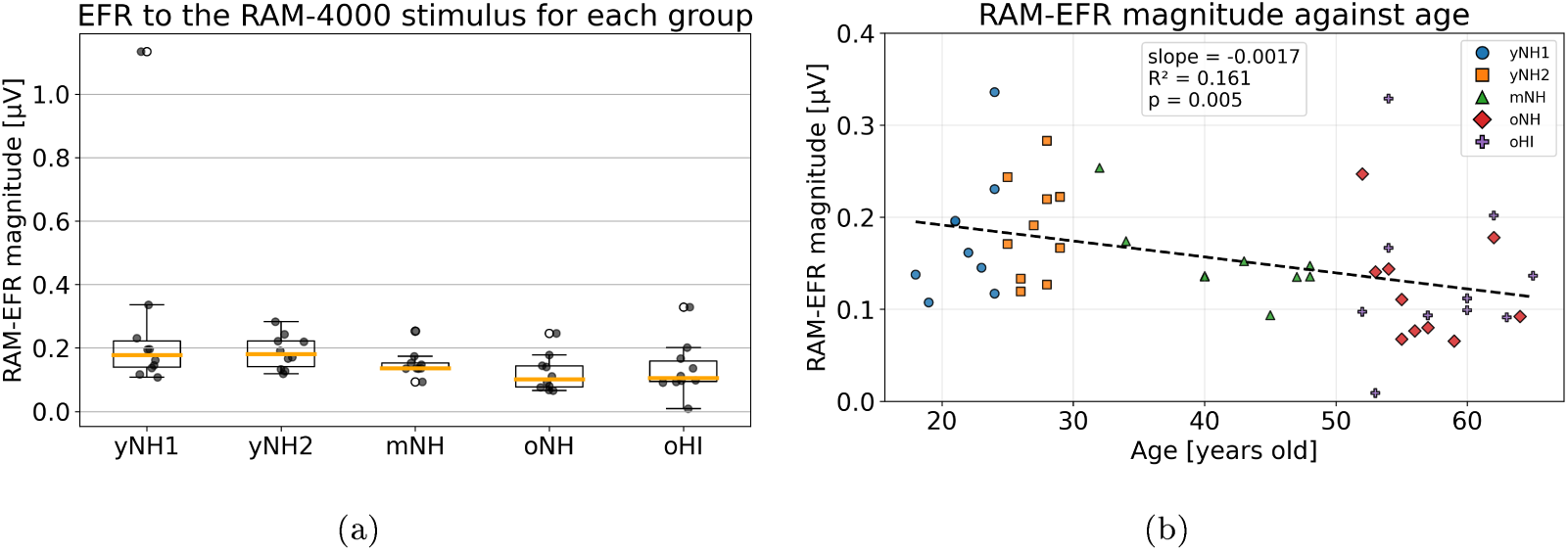
Group comparison of the RAM-EFR magnitude for each group (a) and scatter plot of the RAM-EFR magnitude against age (b). The slope (*β*), and coefficient of determination (*R*^2^) of the linear regression were calculated across all groups, with the associated p-value.

Next, we analyzed the EEG recordings in response to the speech-like phoneme stimulus pairs, differing in spectral regions below the PLL (/du/- /bu/, /o/-/u/ (from /du/), predominantly targeting TFS contrast encoding), or above the PLL (/di/-/bi/, /i/ (from /di/) -/y/, targeting mainly TENV contrast encoding). The resulting EFR frequency spectra for the vowel stimulus pairs are shown for each group in Appendix Figure A1, and the spectra for the syllable pairs are shown in Appendix Figure A2. Figure 7 shows significant within-group differences of the EFR peak magnitude (as in Equation 1) for each stimulus and each group, and Figure 8 shows significant between-group differences in the EFR magnitudes.

**Figure 7:**
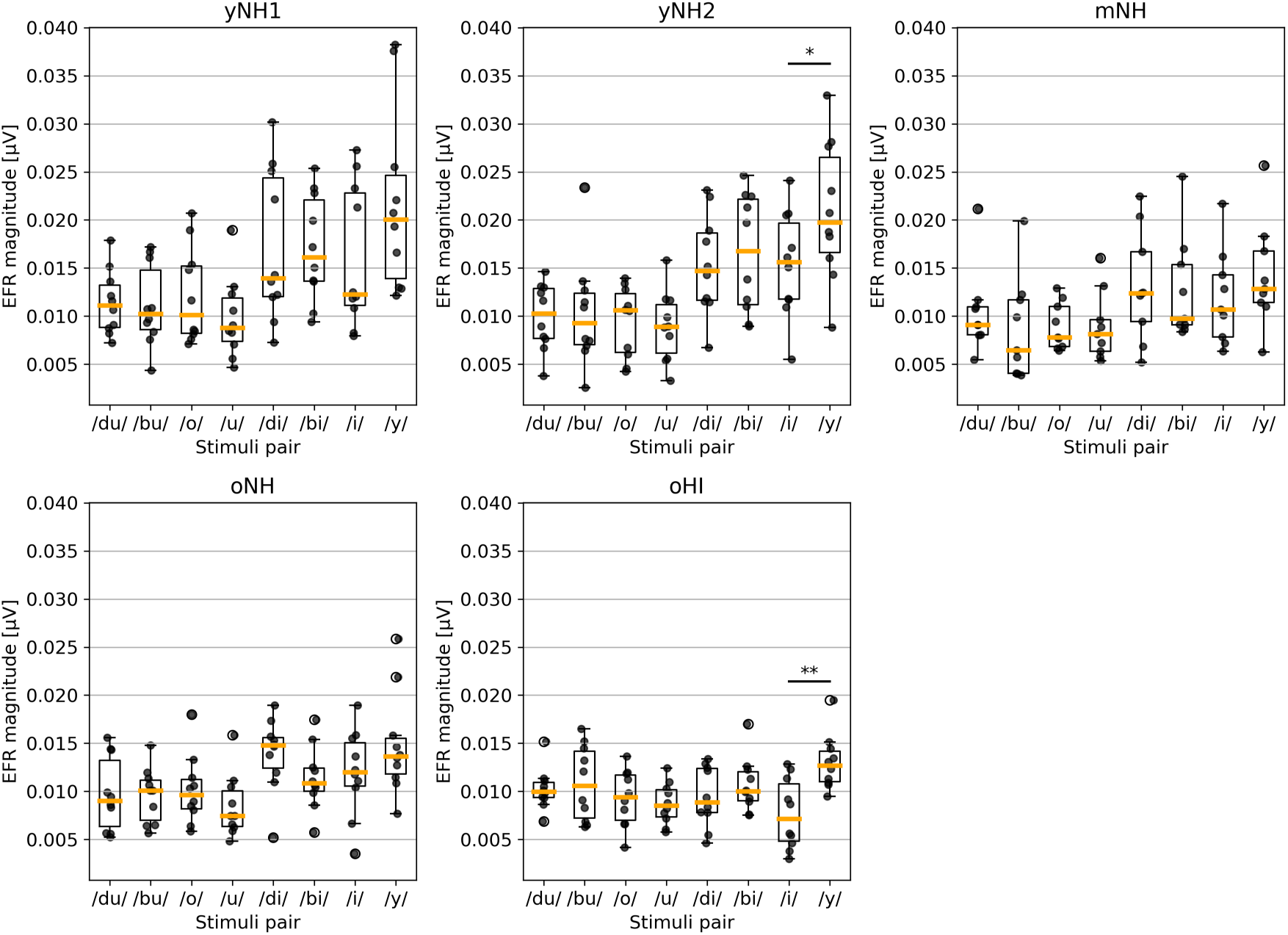
EFR peak magnitude within-group comparisons for each stimulus pair (*p <* .001: * * *; *p <* .01: **; *p <* .05: *).

**Figure 8:**
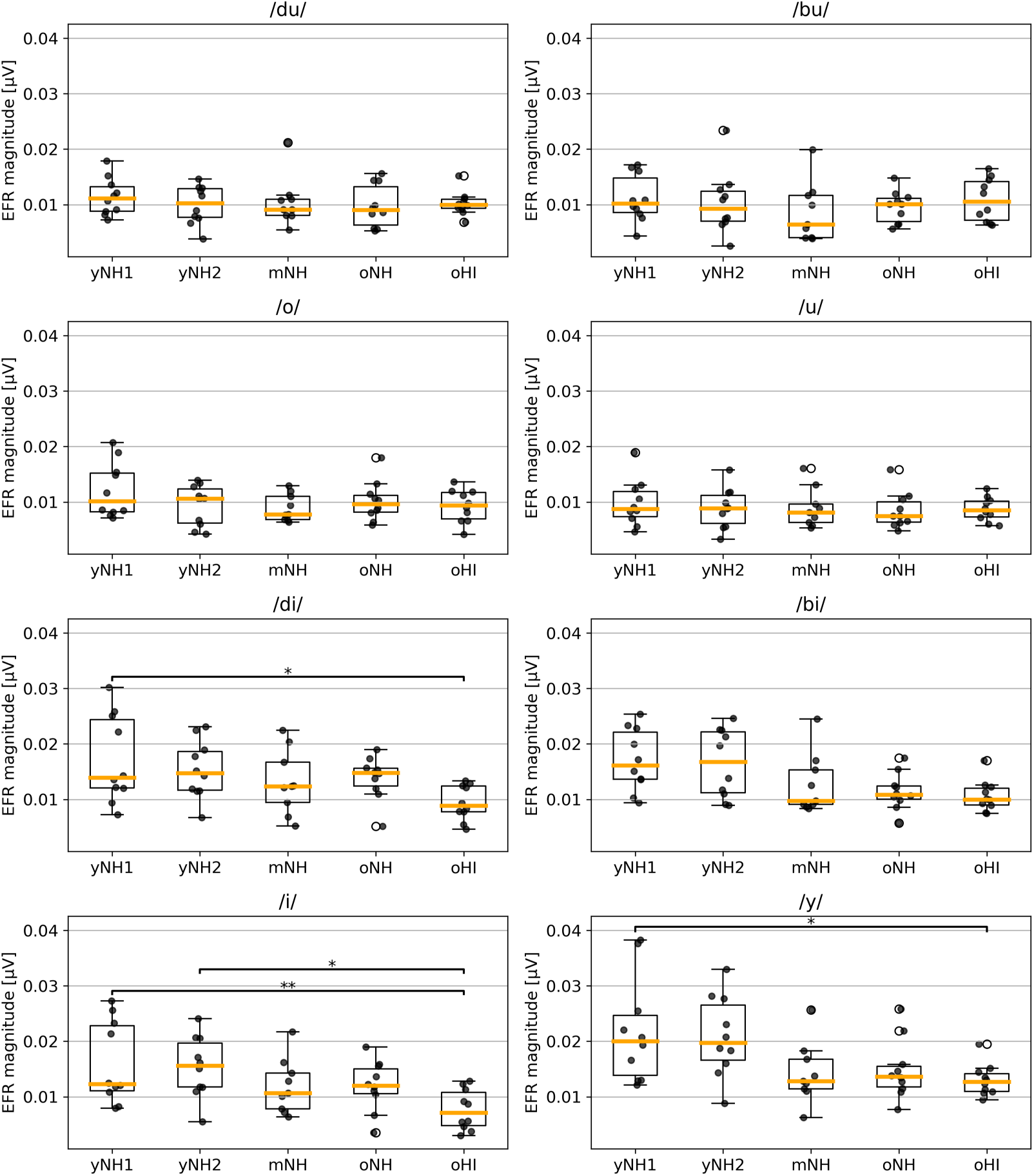
EFR peak magnitude between-group differences for each phoneme (*p <* .001: *\* * *; p <* .01: **; *p <* .05: *).

For the stimulus pair /i/-/y/, the EFR peak magnitude was larger in response to /y/ than to /i/ for each group, but this was only significant for the groups yNH2 *(p <* .05) and oHI *(p <* .01). For groups yNH1, yNH2 and mNH, the harmonics up to the fourth were clearly present in the EFR spectrum for all phoneme pairs (Appendix Figures A1 and A2), while the fourth harmonic was not apparent for group oNH and the third and fourth harmonics were not apparent for group oHI.

The EFR magnitude in response to /i/ was larger for groups yNH1 *(p <* .01) and yNH2 (*p <* .05) than for group oHI. For the stimulus /y/, the EFR response was significantly larger for group yNH1 than for group oHI *(p <* .05). The EFR magnitude to the stimulus /di/ was significantly larger for group yNH1 than for group oHI *(p <* .05). For the stimulus /bi/, the EFR magnitude tended to decrease with age, but this decrease was not significant.

For the stimulus pairs /o/-/u/ and /du/-/bu/, there were no significant between- or within-group differences. These results show that for the stimulus pairs for which the contrast was below the PLL, with most energy at frequencies below the PLL, there was no significant change in the magnitude of the EFR with age. The EFR does not measure TFS fidelity. Because the EFR in response to /o/-/u/ and /du/-/bu/ were largely unaffected by age, our findings suggest that envelope coding derived from lower-frequency cochlear regions is unaffected by age-related neural degradation. Despite the differences between the stimulus pairs below the PLL-specifically, differences in the lower harmonics of *F*_0_ for /o/-/u/ and in the first and second formants around 0.8 - 1.2 kHz for /du/-/bu/-their EFR magnitudes did not differ significantly. However, we observed that the harmonics of *F*_0_ became less apparent with age, as reflected in the EFR frequency spectra presented in Appendix Figures A1 and A2. This decrease in EFR magnitude of the harmonics represents less neural synchrony to the components of the stimulus envelope at the fundamental frequency and its harmonics. Such changes may be related to altered AN encoding of amplitude modulation, potentially involving CS that is selective for high-threshold, LSR AN fibers (Kujawa and Liberman, 2015; Carney, 2018; Furman et al., 2013). It might also be a consequence of reduced phase-locking, particularly at supra-threshold levels.

For the stimulus pairs for which the contrast was above the PLL, and which contained most energy at higher frequencies, there were significant between-group differences in the magnitude of the EFR in response to /i/, /y/ and /di/, consistent with the proposal that TENV processing by high-CF AN fibers declines with age (Dapper et al., 2025). Although CS affects AN fibers across CFs, envelope coding derived from higher-frequency cochlear regions appears to be more vulnerable to age-related neural degradation than envelope coding dominated by lower-frequency regions. This age-related decline in high-CF TENV coding was most pronounced in subjects with elevated PTA4 thresholds. Namely, EFR magnitudes for the young group differed significantly from those of the oHI group, but not of the oNH group (Figure 8). This pattern suggests that age-related CS alone does not fully account for the reduction in high-CF TENV encoding, and that OHC dysfunction further exacerbates the decline. For the stimulus pairs /i/-/y/ and /di/-/bi/, the harmonics of *F*_0_ decreased in magnitude with age, as shown in Appendix Figures A1 and A2.

The effect of age was explored by computing the correlation between the EFR magnitude and age for each stimulus across all groups, as shown in Appendix Figure A4. There was no significant correlation between EFR magnitude and age for the stimuli /du/, /bu/, /o/ and /u/, for which the EFR is generated mainly by TENV coding by low-CF AN fibers. For the stimuli /di/, /bi/, /i/ and /y/, which contain more high-frequency energy, there were significant negative correlations between EFR magnitude and age (/di/: *β =* −0.0001, *R*^2^ = .123, *p =* .014; /bi/: *β =* −0.0002, *R^2^* = .231, *p <* .001; /i/: *β =* −0.0002, *R*^2^ = .186, *p =* .002; /y/: *β =* −0.0002, *R*^2^ = .174, *p* = .003), showing that TENV processing by high-CF AN fibers declines with increasing age.

### 3.2. EFR in the time-domain

We also analyzed the EFR in response to the phoneme stimuli in the time-domain. Figure 9 shows the average of the time-domain waveforms of the EFR for the syllable pairs for each group. Appendix Figure A3 shows the time-domain EFRs for the vowel pairs.

**Figure 9:**
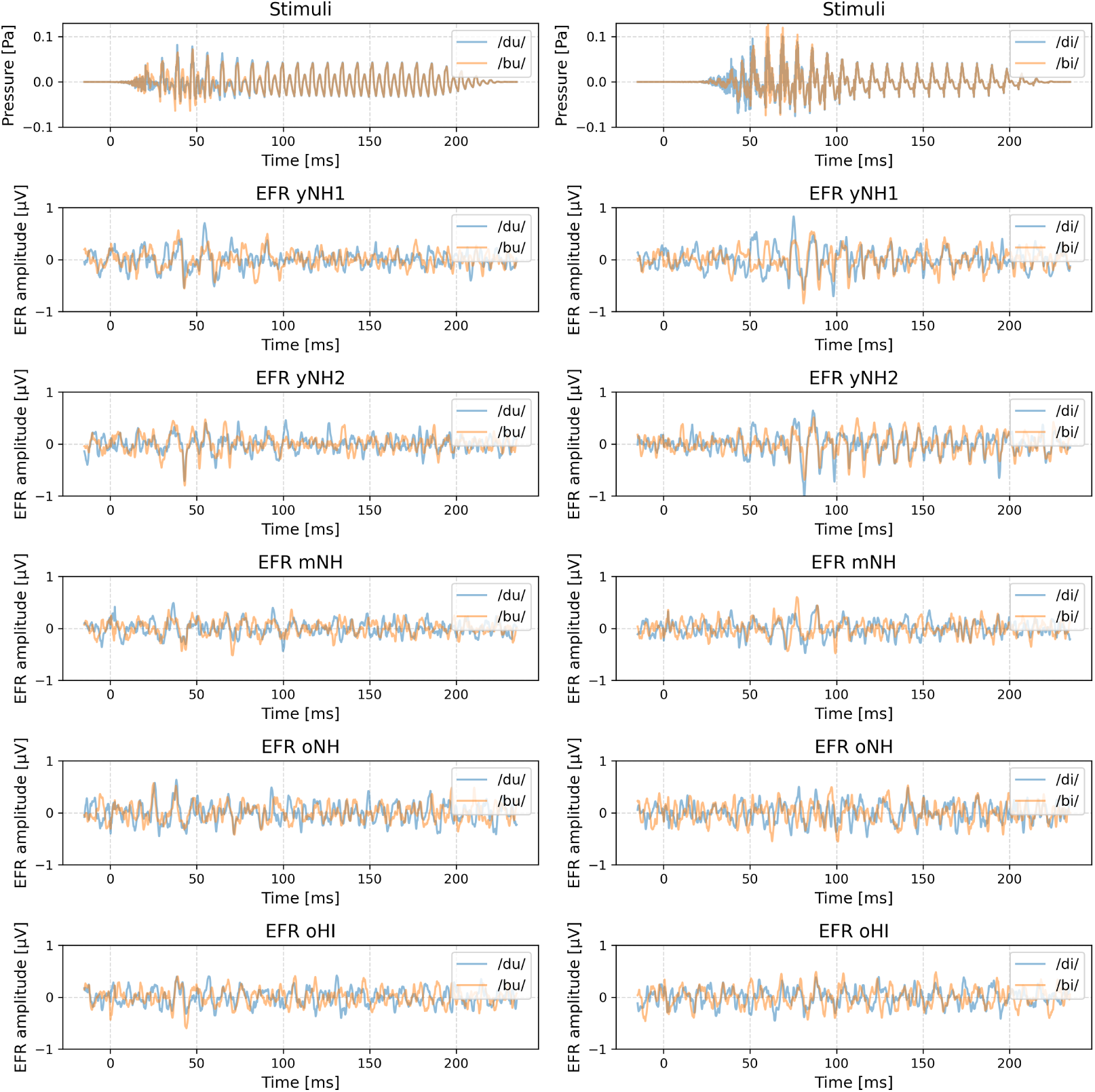
Time-domain waveforms of the EFR, for each syllable pair. Stimulus waveforms are shown in the top panels.

The peaks in the EFR waveform reflect neural phase locking to the TENV of the stimulus. The neural transmission delay is typically between 7 to 9 ms, due to the travel time from the ear to the midbrain (Chandrasekaran and Kraus, 2010). Synchronous firing at stimulus onset mainly depends on high-spontaneous rate (HSR), low-threshold AN fibers, which determine response latencies and perception thresholds (Rhode and Smith, 1986; Meddis, 2006; Heil et al., 2008); the LSR, high-threshold AN fibers contribute little to overall synchrony (Huet et al., 2019). Peaks occurring within the first 50 to 60 ms primarily correspond to the consonant portion of the syllable (/d/ or /b/), which are characterized by acoustic bursts. The later more periodic oscillations correspond to the vowel segment (/u/ or /i/), which has a sustained harmonic structure that drives strong regular neural synchronization.

To investigate whether differences between the stimulus pairs occurred in the EFR time-domain responses, we performed cross-correlation separately for the consonant and vowel segments. Figure 10 shows the cross-correlation values for each syllable pair and each group. For the pair /du/-/bu/, the cross-correlation values for the consonant segments showed a significant group effect. Although the data for most groups passed the Shapiro-Wilk normality test, one group did not (oHI: *W* = .8314, *p* = .0348), prompting the use of the Kruskal-Wallis H test, which revealed a significant difference across groups *(H* = 12.3889, *p* = .0147). Post hoc Dunn’s tests indicated marginally significant differences between groups oHI and yNH2 (*p* = .0440) and between groups oHI and mNH *(p* = .0440), suggesting reduced neural synchrony in the oHI group. For the vowel segments of the same pair, normality was violated for group oNH *(W* = .7609, *p* = .0048), and the Kruskal-Wallis test did not reach significance *(H* = 8.1161, *p* = .0874), indicating no strong group-level differences.

**Figure 10:**
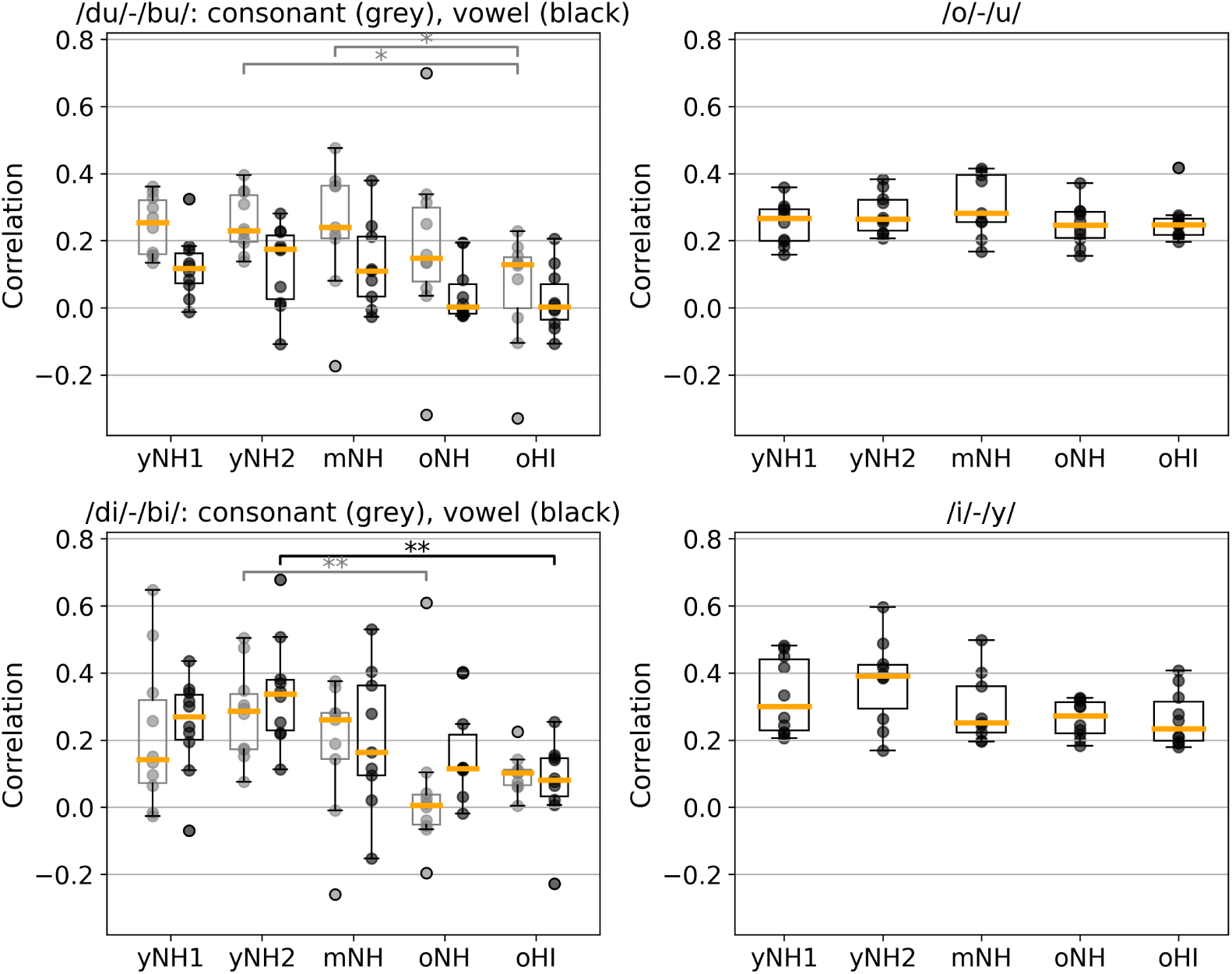
Cross-correlation values for each phoneme pair and each group (*p <* .001: ***; *p <* .01: **; *p <* .05: *). Correlation values for consonant and vowel sections are shown in grey and black, respectively.

For the pair /di/-/bi/, the consonant segment analysis showed a violation of normality for group oNH *(W* = .7251, *p* = .0018), and the Kruskal-Wallis test was significant (*H* = 14.9256, *p* = .0049). Dunn’s test revealed a significant difference between groups oNH and yNH2 *(p* = .0036), indicating altered consonant processing for the oNH group. For the vowel segment, normality was violated for the oNH group *(W* = .8443, *p* = .0497), and the Kruskal-Wallis test was again significant (*H* = 12.3010, *p* = .0152). Dunn’s test showed a significant difference between groups oHI and yNH2 *(p* = .0096), suggesting reduced vowel similarity for the oHI group.

For the vowel-only pair /o/-/u/, the data for group oHI did not pass the normality test *(W* = .7676, *p* = .0059), and the Kruskal-Wallis test was not significant (*H* = 3.5579, *p* = .4691), suggesting comparable vowel segment similarity across groups. For the vowel-only pair /i/-/y/, all groups passed the normality test, and the ANOVA did not reveal significant differences (*F* = 2.3234, *p* = .0714). Across all five subject groups, the cross-correlation between the EFR time-domain waveforms evoked by the two stimuli within each vowel pair was low (approximately 0.2 - 0.4), indicating weak similarity and no meaningful correlation between the responses.

In summary, EFR time-domain waveforms for the consonant segments showed that yNH subjects exhibited higher correlations than older and HI groups, for stimulus pairs with contrast both below the PLL (/du/-/bu/) and above (/di/-/bi/). This suggests that younger subjects produce more stable and consistent neural responses across similar consonant contexts, whereas older and HI subjects show greater neural variability, leading to lower correlation values. Importantly, higher response stability does not on its own guarantee better behavioral discrimination. While the correlation metric captures trial-to-trial consistency of the neural response, perceptual decisions additionally depend on how distinct the neural representations of the two stimuli are. In other words, behavioral performance reflects a combination of both factors: the reliability with which each stimulus is encoded across trials and the degree to which their neural representations can be discriminated from one another. Thus, increased encoding fidelity may coexist with comparable discrimination performance. Vowel-only pairs (/o/-/u/, /i/-/y/) showed no significant correlations, and no significant group differences, confirming that age-related changes primarily affect the encoding of rapid consonant cues rather than sustained vowel periodicity.

### 3.3. Phoneme discrimination

Figure 11 shows phoneme discrimination scores for each group and each noise condition and stimulus pair. For the pair /du/-/bu/, the score with ipsilateral noise was significantly higher than in quiet for groups yNH1 (*p <* .05), yNH2 *(p <* .001) and mNH (*p <* .001). For groups oNH and oHI, discrimination with ipsilateral noise was also better than in quiet, although this difference did not reach statistical significance. For groups yNH2 *(p <* .001) and mNH *(p <* .001), the score was higher with ipsilateral noise than with contralateral noise. There were no significant between-group differences for the /du/-/bu/ pair. In quiet and with contralateral noise, discrimination for this pair was close to chance level for all groups, suggesting that this stimulus contrast was the most difficult to discriminate, indicating weak or ambiguous cues. Ipsilateral noise typically degrades speech perception, therefore, the improvement observed for the /du/-/bu/ contrast may suggest that the masker altered the relative salience of auditory cues. Ipsilateral noise may have suppressed misleading information or shifted cue weighting, thereby enhancing the effective consonant contrast. Although counter-intuitive, improvements with added ipsilateral noise have been documented. One well-known example is the phenomenon of “phonemic restoration” (Warren, 1970; Başkent, 2010), where added noise can improve intelligibility. This effect is typically observed with stimuli containing artificial interruptions that introduce misleading phonetic cues; filling these gaps with noise masks the misleading information and facilitates perceptual restoration (Bhargava et al., 2014). Although our stimuli did not include interruptions, their chimeric construction introduced unnatural cue combinations. In particular, for the /du/-/bu/ contrast, acoustic differences that are normally present in the high-frequency region were neutralised in the present design. Younger subjects may expect these cues in quiet and be disrupted by their absence, whereas in noise they may adopt a different strategy and attribute missing information to masking rather than to the signal itself. Although this complicates interpretation of our results, it also suggests that participants processed the stimuli as speech, supporting our goal of probing speech-specific perception.

**Figure 11:**
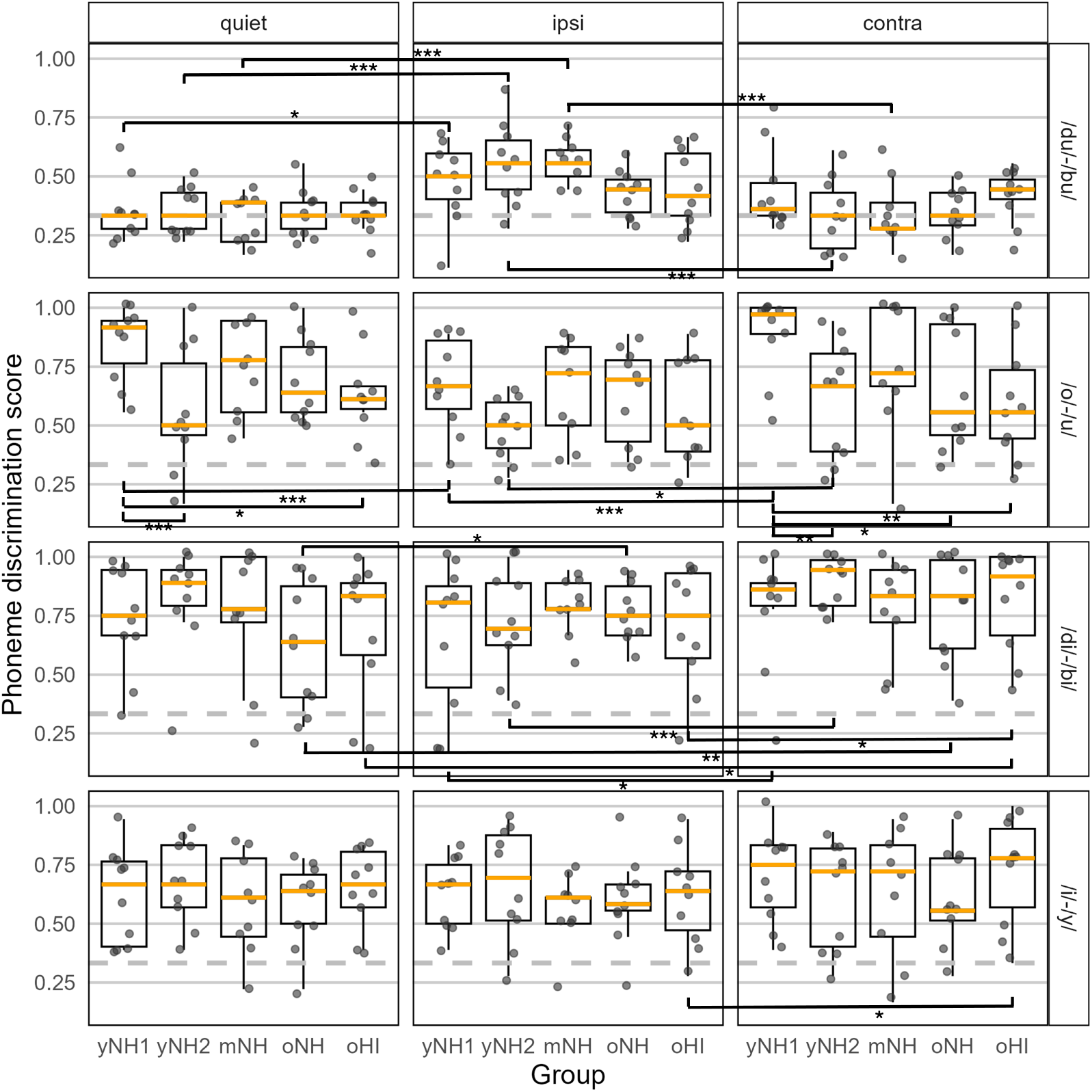
Phoneme discrimination scores for each group, stimulus pair and noise condition (*p <* .001: * * *; *p* < .01: **; *p <* .05: *).

For the stimulus pair /o/-/u/, the discrimination score for group yNHl was significantly lower with ipsilateral noise than in quiet *(p <* .001) and with contralateral noise (*p <* .001). For group yNH2, the score with ipsilateral noise was also significantly lower than with contralateral noise *(p <* .05). In quiet, the score for group yNHl was significantly higher than for groups yNH2 *(p <* .001) and oHI (*p <* .05). The observed decline in /o/-/u/ dis-crimination -a contrast thought to rely relatively strongly on TFS cues-with increasing age and hearing impairment in quiet, is consistent with findings of Schirmer et al. (2024). With contralateral noise, the score for group yNHl was significantly higher than for groups yNH2 *(p <* .01), oNH *(p <* .05) and oHI (*p <* .01). With ipsilateral noise, the discrimination scores were similar for all groups.

For the stimulus pair /di/-/bi/, all groups performed similarly for each noise condition, but there were significant effects of noise type. For the yNH1 group, the score was significantly higher with contralateral noise than with ipsilateral noise *(p <* .05), which was also the case for the yNH2 group *(p <* .01) and oHI group *(p <* .05). For the oNH group, the score in quiet was significantly lower than with ipsilateral noise *(p <* .05). For the oNH *(p <* .01) and oHI *(p <* .05) groups, the score with contralateral noise was significantly higher than in quiet, which may be a consequence of altered listening strategy or cue weighting with contralateral noise in older subjects, for example by biasing attention toward the ear receiving the stimulus pair (Wendt et al., 2021). Moreover, the phenomenon of “phonemic restoration” due to the chimeric construction of the /di/-/bi/ stimulus pair, could potentially explain why the performance for the older groups was better with noise than in quiet, since these subjects may have developed extra reliance in quiet on a low-frequency cue that has been removed in our design (Warren, 1970; Başkent, 2010; Bhargava et al., 2014). In the young groups, no such improvement with contralateral noise relative to quiet was observed, likely because their performance in both quiet and contralateral noise was already at “ceiling” for the /di/-/bi/ contrast, leaving little room for further enhancement.

Finally, for the stimulus pair /i/-/y/, the score for the oHI group was significantly higher with contralateral noise than with ipsilateral noise *(p <* .05), as found by Schirmer et al. (2024). Apart from this difference, the noise condition had no influence on the score for any of the groups.

## 4. Discussion

The present study examined how age-related CS and OHC damage affect peripheral speech encoding and phoneme discrimination, using electrophysiological markers (RAM-EFR and speech-evoked EFRs) and behavioral discrimination tasks in several noise conditions. Our results demonstrate distinct effects of aging, CS, and OHC dysfunction on electrophysiological and behavioral measures that are thought to reflect the encoding of TENV and TFS cues, extending the recent findings of Schirmer et al. (2024) and Dapper et al. (2025) on cortical and midbrain correlates of degraded auditory encoding.

### 4.1. Age-related EHF degradation and RAM-EFR sensitivity to CS

The progressive elevation in PTA-EHF thresholds in the mNH group and older groups confirms early high-frequency degradation with age. Although elevated audiometric thresholds primarily reflect OHC dysfunction rather than synaptic loss, histopathological and physiological evidence indicates that EHF regions of the cochlea are particularly vulnerable to age-related damage (Liberman et al., 2016; Bramhall et al., 2017). In this basal region, OHC dysfunction and neural degeneration related to CS may therefore occur in parallel. As also observed by Schirmer et al. (2024), RAM-EFR magnitudes significantly decreased with age, while no significant differences were observed between oNH and oHI groups, consistent with evidence that RAM-EFRs primarily reflect age-related CS, while being relatively insensitive to audiometric threshold differences (Keshishzadeh et al., 2020, 2021; Van Der Biest et al., 2023; Vasilkov et al., 2021). Together, these findings support the view that CS begins around midlife, prior to measurable hearing loss at conventional audiometric frequencies (e.g. PTA4), and progresses alongside EHF threshold elevation.

### 4.2. EFR in the frequency domain: high-CF versus low-CF TENV coding

EFR magnitudes in response to phoneme stimuli, as shown in Figure 8, differed across stimulus pairs and subject groups. Stimuli containing most energy below the PLL (/du/-/bu/, /o/-/u/), for which the EFR is mainly determined by low-CF TENV coding of the AN fibers, showed preserved EFR magnitudes in quiet across age groups, whereas stimuli containing most energy above the PLL (/di/(-/bi/), /i/-/y/), for which the EFR primarily depends on TENV encoding by high-CF fibers, exhibited significant group differences, particularly between yNH and oHI subjects. A decline with age in EFR magnitude in response to phonemes relying on high-CF TENV coding was also observed by Dapper et al. (2025). This pattern indicates that high-frequency TENV processing is more susceptible to age and hearing impairment than low-frequency TENV coding, consistent with physiological and modeling studies showing a decline of TENV phase-locking at higher CFs with increasing age or severity of SNHL (Henry and Heinz, 2012; Parthasarathy and Kujawa, 2018).

The finding that only the oHI group, characterized by both CS and OHC damage, showed robust reductions in EFR magnitude above the PLL suggests that OHC dysfunction exacerbates age-related declines in high-CF envelope encoding. Although reduced cochlear compression associated with OHC loss can increase suprathreshold envelope modulation depth and thereby enhance TENV representation (Vasilkov and Verhulst, 2019; Dreyer and Delgutte, 2006), this potential benefit may only be exploited when neural synchrony is sufficiently preserved. In the presence of age-related CS, however, the auditory nerve may be unable to encode the increased envelope contrast effectively, leading to a net reduction in EFR magnitude (Verhulst et al., 2016). The absence of a significant RAM-EFR difference between the oNH and oHI groups indicates that both groups likely exhibited a similar degree of age-related CS. The additional reduction in high-CF TENV coding observed only in the oHI group therefore reflects the interaction between CS and OHC dysfunction: when synaptic integrity is already compromised, the auditory system might not be able to take advantage of the increased modulation depth caused by OHC loss, resulting in a net degradation of envelope encoding.

While RAM-EFR magnitudes were relatively insensitive to OHC loss, speech-evoked EFRs targeting high-CF TENV coding appear to capture combined degradations of neural synchrony and cochlear mechanics. This interpretation is consistent with recent work demonstrating that while RAM-EFR paradigms are optimized to isolate CS and are relatively insensitive to OHC loss, EFRs elicited by more complex speech-like stimuli reflect broader cochlear mechanics and neural synchrony, making them vulnerable to both pathologies (Van Der Biest et al., 2023; Keshishzadeh et al., 2020; Vasilkov et al., 2021).

### 4.3. EFR in the time-domain: neural synchrony

Analysis of the EFR time-domain waveforms showed that cross-correlations between responses to phoneme pairs were highest for yNH subjects and significantly reduced for oNH and oHI groups for consonant segments, as shown in Figure 10. These findings suggest increased neural variability and reduced temporal precision in envelope coding in older subjects. Such changes have previously been linked to degraded AN encoding, including reduced synchrony to stimulus envelopes and increased temporal jitter in animal and human studies of aging (Furman et al., 2013; Sergeyenko et al., 2013). The fact that vowel-only segments showed smaller age effects reinforces the interpretation that TENV encoding of consonants, which depends heavily on accurate envelope onset timing, is particularly vulnerable to CS (Bharadwaj et al., 2015).

### 4.4. Phoneme discrimination and the role of noise configuration

Behavioral phoneme discrimination revealed interactions between stimulus type, age, and noise configuration, as shown in Figure 11. For the /du/-/bu/ pair, which is thought to rely predominantly on TFS-related cues, yNH and mNH subjects showed significantly better discrimination with ipsilateral noise than in quiet or with contralateral noise. No such effect was observed in the older groups (oNH and oHI), whose performance was close to chance level, limiting the interpretation of noise-related differences in these groups. In younger subjects, the improvement with ipsilateral noise may reflect more intact TFS processing compared to older subjects, which could facilitate cue segregation when target and masker are co-located (Parthasarathy and Kujawa, 2018; Plack et al., 2014). As noted previously, this pattern was unexpected, although improvements with ipsilateral noise have been reported previously (Warren, 1970; Başkent, 2010; Bhargava et al., 2014). In the present case, the chimeric stimuli may have introduced atypical cue combinations, and for the /du/-/bu/ contrast in particular, high-frequency cues were neutralised. This may have disrupted subjects in quiet but led them to adopt a different cue-weighting strategy in noise. Thus, the addition of ipsilateral noise may have altered the relative salience of available cues, although the present data do not allow the underlying mechanism to be identified.

For the /o/-/u/ pair, which also predominantly targets TFS coding but without a consonant segment, the pattern was reversed: discrimination with ipsilateral noise was worse than in quiet and with contralateral noise for yNH subjects, whereas older groups showed no significant effect of noise condition. Younger subjects rely strongly on low-frequency TFS cues, which are masked when noise is presented to the same ear, whereas older subjects, who are already affected by CS, have reduced access to TFS cues and thus show less change across conditions (Kortlang et al., 2016; Füllgrabe, 2013). There were significant between-group differences for the discrimination of the /o/-/u/ stimulus pair. In quiet, the yNH group performed significantly better than the oHI group, but not than the oNH group, suggesting that hearing impairment in addition to age is associated with reduced performance for this TFS-based contrast. With contralateral noise, the yNH group performed significantly better than all older subjects. This pattern suggests that age-related changes in binaural or central auditory processing, which are particularly engaged by contralateral masking, adversely affect discrimination performance for cues that are thought to rely predominantly on TFS processing (Wendt et al., 2021).

For the /di/-/bi/ stimulus pair, which primarily targets TENV processing, the older groups exhibited poorer consonant discrimination in quiet and with ipsilateral noise than with contralateral noise. Because contralateral noise does not directly mask the target signal at the stimulated ear, the mechanism underlying this finding remains unclear. One possible explanation is that contralateral noise may alter listening strategy, cue weighting, or cause better concentration on the ear in which the stimulus pair was presented in older subjects (Wendt et al., 2021). However, the present data do not allow us to distinguish between these potential mechanisms, and these interpretations therefore remain speculative. No improvement with contralateral noise relative to quiet was observed for the yNH groups, possibly due to a “ceiling” effect for this stimulus pair in quiet and with contralateral noise. For the /i/-/y/ vowel discrimination, which also relies mainly on TENV coding, scores were lower with ipsilateral noise than with contralateral noise for the oHI subjects. This agrees with the expectation that speech-in-noise understanding depends on intact TENV encoding, and that degradation of high-CF envelope coding disproportionately affects performance in conditions where peripheral envelope cues are disrupted, such as in the presence of modulation masking from ipsilateral noise (Henry and Heinz, 2012; Oxenham, 2016; Stone et al., 2011, 2012; Stone and Moore, 2014).

### 4.5. Relation of phoneme discrimination to EFR magnitude

The effects of age and hearing loss on the EFR magnitude and performance of the phoneme discrimination task remain puzzling. On one hand, stimuli whose discrimination predominantly relies on TENV coding (/di/-/bi/, /i/-/y/) showed significantly reduced EFR representation for oHI subjects compared to younger subjects (Figure 8). On the other hand, this reduced EFR magnitude was not associated with poorer behavioral discrimination; performance was similar across age groups (Figure 11). These findings raise the possibility that oHI subjects compensate for degraded envelope encoding through alternative acoustic cues or central mechanisms such as increased attentional control or auditory gain (Auerbach et al., 2014; Pichora-Fuller et al., 2016; Peelle, 2018; Goossens et al., 2018; Parthasarathy et al., 2019). EFRs for stimulus pairs predominantly targeting TFS contrast coding (/o/-/u/) remained stable across age and hearing status (Figure 8), yet psychophysical performance changed significantly in quiet and with contralateral noise (Figure 11).

The /i/-/y/ stimuli differ in high-frequency formant frequencies (2-4 kHz). Behavioral discrimination of this contrast is therefore expected to rely mainly on place-based spectral cues, but TFS cues may also contribute, particularly for listeners with intact temporal coding. Because place-based spectral cues are generally more robust to aging and SNHL than TFS-based cues, discrimination of the /i/-/y/ contrast is expected to be less strongly affected by age-related declines in temporal precision than discrimination of TFS-based contrasts, such as the /o/-/u/ pair (Whiteford et al., 2017). Based on the age-related increase in PTA-EHF thresholds shown in Figure 2b, indicating peripheral high-frequency cochlear degradation, we would expect the envelope tracking ability in response to /i/-/y/ to decline with age, which would be consistent with the observed decline of the EFR measured by EEG.

The /o/-/u/ stimuli differ in the lower harmonics of *F*_0_ (116 Hz). Discrimination of this low-frequency contrast relies more on low-frequency TFS coding, which we expect to degrade with age (Ponsot et al., 2025; Moore et al., 2012) and may be related to poorer behavioral discrimination. However, the EFR predominantly reflects an envelope-following measure and does not measure TFS fidelity directly. Therefore, the /o/-/u/ EFR magnitude was almost unchanged with age, since low-CF fibers responsible for the envelope tracking are not as much affected by age as high-CF fibers. This disconnect between TFS and TENV processing in the EFR might explain why, for the /o/-/u/ contrast, the EFR responses were unaffected by age while discrimination scores did show effects of age and SNHL on TFS discrimination. Together, the prominent role of place-based spectral cues, alongside possible contributions from TFS cues, for /i/-/y/ discrimination, and the stronger reliance on TFS cues for /o/-/u/ discrimination, plus the TENV-dominant nature of the EFR, may explain why, with age and SNHL, /i/-/y/ showed reduced EFR magnitude without a corresponding behavioral loss, whereas /o/-/u/ showed stable EFRs with behavioral loss.

The dissociation between neural and behavioral measures observed here closely parallels the dissociation reported by Ponsot et al. (2025), who demonstrated robust age- and hearing loss-related reductions in neural phase-locking to both TENV components, assessed using a high-frequency (4 kHz) RAM tone, and TFS components, assessed using a low-frequency (< 1.5 kHz) spectrally modulated complex tone, yet found no relationship between these EFR markers and psychophysical performance. Furthermore, Goossens et al. (2018) demonstrated that stronger neural envelope encoding, measured at both subcortical and cortical levels, can paradoxically be associated with poorer speech perception in noise for both NH and HI adults. This finding supports our interpretation that changes in neural synchrony, as reflected by EFR magnitude, do not necessarily translate into functional perceptual outcomes, consistent with the dissociation between neural and behavioral measures observed in the present study.

The mismatch between electrophysiological and psychophysical outcomes suggests that behavioral performance is not determined solely by the fidelity of peripheral neural encoding (Parthasarathy and Kujawa, 2018; Plack et al., 2014). Instead, perception likely reflects an interaction between sensory encoding limits on the neural representation itself (e.g. reduced phase-locking or neural synchrony) and perceptual strategies, such as differential weighting of available acoustic cues. These unresolved discrepancies highlight the need for further investigation into how neural coding deficits are related to perceptual outcomes, and whether redundancy of cues (Plack et al., 2014) or top-down processes (e.g. attentional modulation, listening effort, or context-dependent cue weighting) mitigate the impact of degraded TENV or TFS encoding (Pichora-Fuller et al., 2016).

### 4.6. Limitations and future directions

While the present study provides valuable insights into how aging, CS and OHC damage jointly shape peripheral speech encoding, several limitations should be acknowledged. The group sizes were modest, which may have limited the statistical power to detect subtle between-group differences in electrophysiological measures. Interindividual variability in EFR magnitude, likely influenced by anatomical and recording factors, such as electrode placement variability, may also have masked small group differences (Bidelman et al., 2026; Coffey et al., 2019; McFarlane and Sanchez, 2023). Another limitation of this study is the absence of detailed noise-exposure histories, which are relevant for interpreting CS-related effects. While subjective hearing evaluations were collected, they do not capture cumulative exposure.

It is also worth noting that behavioral discrimination performance was at chance for most subjects for some conditions (/du/-/bu/), or was at “ceiling” (/di/-/bi/), which might have masked a more complex picture. Further studies may help clarify these effects by better adjusting the difficulty level of discrimination for each phoneme pair. Furthermore, while RAM-EFRs and speech-evoked EFRs provide complementary insights into subcortical temporal processing, they cannot fully isolate peripheral from central contributions. Future work could better disentangle cortical from subcortical generators by systematically varying modulation rates, as higher modulation rates (e.g. 80 - 100 Hz) preferentially evoke brainstem-dominated phase-locking, whereas lower modulation rates (e.g. < 40 Hz) elicit stronger cortical phase-locking (Coffey et al., 2016, 2017; Bidelman, 2018; Herdman et al., 2002). Moreover, combining EFRs with source-informed analyses (e.g. exploiting differences in response latency, scalp topography, or source modeling to separate early subcortical from later cortical components) or with complementary cortical measures (e.g. low-frequency cortical phase-locking or late auditory evoked potentials, which predominantly reflect cortical sound processing) could allow a more refined separation of contributions from different levels of the auditory pathway. In addition, future work should further investigate how neural coding deficits are related to perceptual outcomes, and under which conditions neural measures reliably predict behavioral outcomes.

## 5. Conclusions

This study investigated how aging and hearing impairment affect peripheral speech encoding and phoneme discrimination, with a particular focus on CS and OHC damage, across five subject groups differing in age and hearing impairment. RAM-EFR magnitudes were significantly correlated with age, suggesting an age-related decline consistent with progressive CS beginning in midlife (Van Der Biest et al., 2023; Vasilkov et al., 2021). This trend co-occurred with elevation of EHF audiometric thresholds from midlife onward, which primarily reflect OHC dysfunction, but are also known to involve cochlear regions that are particularly vulnerable to neural degeneration related to CS (Bramhall et al., 2017; Lobarinas et al., 2013; Liberman et al., 2016; Wang et al., 2021; Wu et al., 2019). Together, these findings support the view that CS may begin around midlife, prior to measurable hearing loss at conventional audiometric frequencies (e.g. PTA4), and may co-occur with EHF threshold elevation (Bramhall et al., 2017; Lobarinas et al., 2013; Liberman et al., 2016).

Speech-evoked EFRs recorded in quiet provided insight into the specific coding mechanisms affected. Stimuli containing most energy below the PLL, for which the EFR is predominantly determined by low-CF TENV contributions, showed largely preserved neural encoding across age groups. In contrast, stimuli containing most energy above the PLL, for which the EFR is predominantly determined by high-CF TENV contributions, exhibited reduced EFR magnitudes with age, especially for the oHI group. These results suggest that high-CF envelope encoding is vulnerable to the combined effects of CS and OHC damage, whereas low-CF envelope coding is comparatively robust. This pattern aligns with findings from Dapper et al. (2025) and Schirmer et al. (2024), who observed that age-related reductions in cochlear synchrony and midbrain onset coding primarily affect TENV contrast representations.

Analysis of the EFR time-domain waveforms supported this interpretation: cross-correlations between responses to stimulus pairs were highest for young subjects and declined with age, particularly for consonant segments, indicating increased neural noise and reduced temporal precision in TENV coding for older subjects. This reduced response consistency is consistent with altered AN population statistics observed in animal studies of noise-induced CS, suggesting a selective loss of LSR and medium-SR high-threshold fibers, which has been identified as a key physiological hallmark of CS (Furman et al., 2013; Liberman et al., 2015). Importantly, selective loss of these fibers can occur without elevated audiometric thresholds at conventional frequencies, while nonetheless degrading neural population synchrony and contributing to difficulties with speech perception in noisy environments (Sergeyenko et al., 2013).

Behavioral phoneme discrimination did not mirror the electrophysiological findings, indicating that behavioral performance is not determined solely by the fidelity of peripheral neural encoding (Parthasarathy and Kujawa, 2018; Plack et al., 2014; Ponsot et al., 2025; Goossens et al., 2018). While EFRs in response to phonemes targeting high-CF fiber envelope coding declined significantly with age, EFRs in response to phonemes targeting low-CF fiber envelope coding remained relatively stable. However, discrimination of phonemes predominantly involving TENV-based contrasts did not show significant effects of age or hearing impairment for any of the noise conditions, while discrimination of phonemes with predominantly TFS-based contrasts showed behavioral decline with age. This pattern can be understood by noting that high-frequency contrasts, above the PLL, are discriminated primarily through place-based spectral cues, which remain robust with age, whereas low-frequency contrasts, below the PLL, rely on TFS coding, a mechanism known to degrade with aging. The EFR predominantly reflects TENV coding, and because age-related neural decline is most pronounced in high-CF regions, EFRs in response to high-frequency contrasts show the largest age-related reductions even though their perceptual discrimination is preserved.

Taken together, these results demonstrate that age-related CS emerges before measurable audiometric loss at conventional audiometric frequencies, progressively degrading high-CF TENV encoding at the brainstem level while leaving low-CF TENV encoding relatively intact. Behavioral discrimination of contrasts above the PLL remained stable with age and hearing impairment, likely because discrimination of the high-frequency contrast in the formants predominantly uses place-based spectral cues, while discrimination of contrasts below the PLL, targeting TFS coding, worsened with age. Integrating electrophysiological measures such as RAM- and phoneme-evoked EFRs with behavioral phoneme discrimination may provide a framework for identifying and characterizing subclinical hearing deficits. Future work should further investigate the link between neural coding deficits and perceptual outcomes, using approaches to better dissociate subcortical temporal processing from peripheral and central contributions. These complementary approaches may help characterize early neural changes beyond conventional audiometric measures, with potential relevance for diagnosis and monitoring of neural auditory decline in aging and HI populations.

## Author declarations

### Author contributions

Conceptualization: S.V., E.G., D.B., L.R. and M.K.; Data curation: J.S. and K.D.; Methodology: M.W., K.D., J.S., E.G., D.B. and S.V.; Formal Analysis: M.W., K.D. and J.S.; Writing - original draft: M.W.; Writing - review and editing: M.W., L.R., M.K., E.G., D.B. and S.V.; Supervision: S.V. All authors have read and agreed to the published version of the manuscript.

### Data availability

The raw data supporting the conclusions of this article will be made available by the authors on request.

## Acknowledgments

This work was supported by the Deutsche Forschungsgemeinschaft DFG KN 316/13-1, DFG RU 713/6-1, KL 1093/12-1; ERA-NET NEURON JTC 2020: BMBF 01EW2102 CoSySpeech and FWO G0H6420N; IZKF Promotionskolleg of the Faculty of Medicine, University Hospital of Tübingen. VICI Grant (Grant No. 918-17-603), Netherlands Organization for Scientific Research (NWO) and the Netherlands Organization for Health Research and Development (ZonMw). The funding sources were not involved in study design; data collection, analysis and interpretation, or writing and submitting the manuscript.

## Appendix

**Figure A1:**
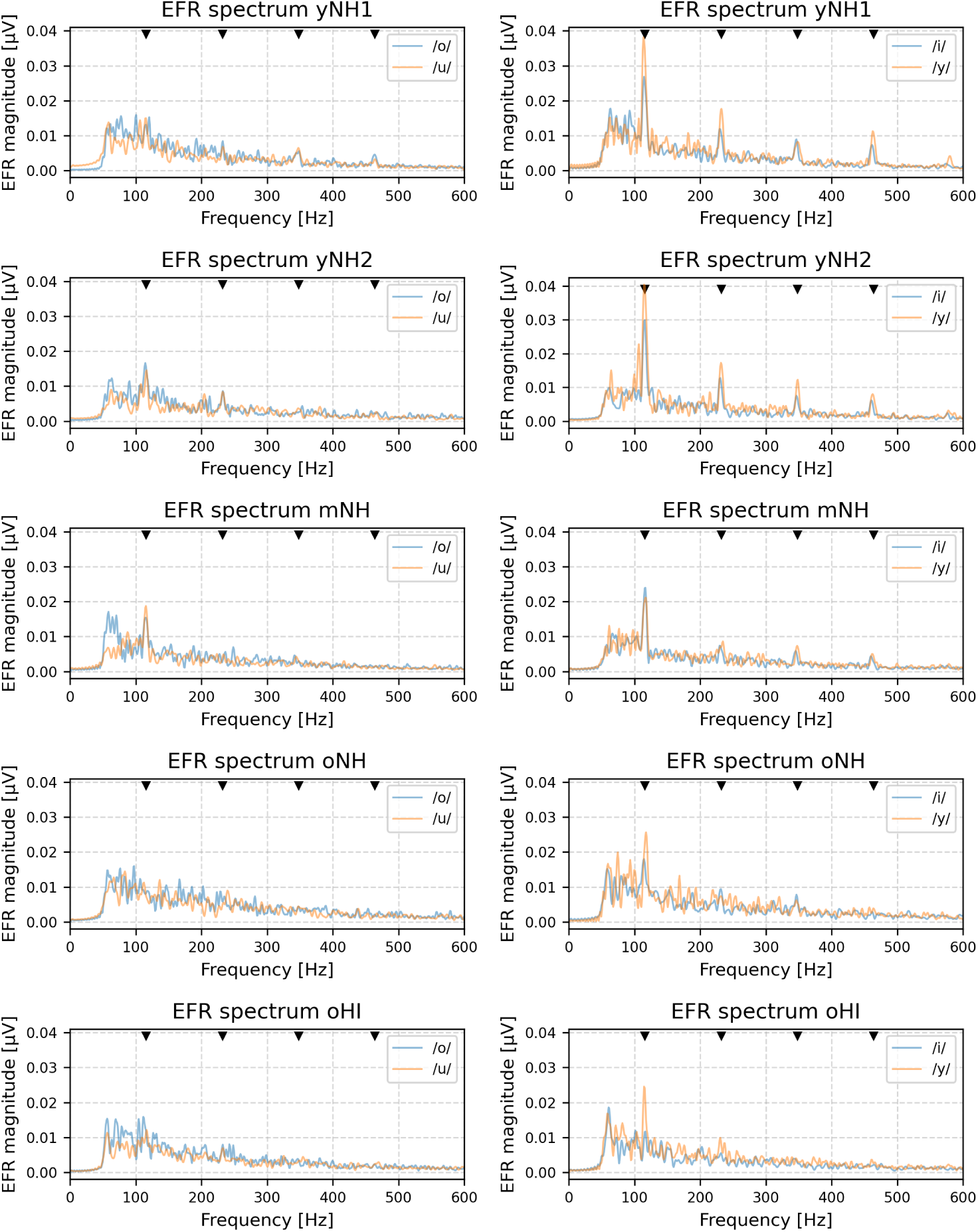
EFR frequency spectra for vowel contrasts for each group. The harmonics of *F*_0_ are marked by black triangle symbols.

**Figure A2:**
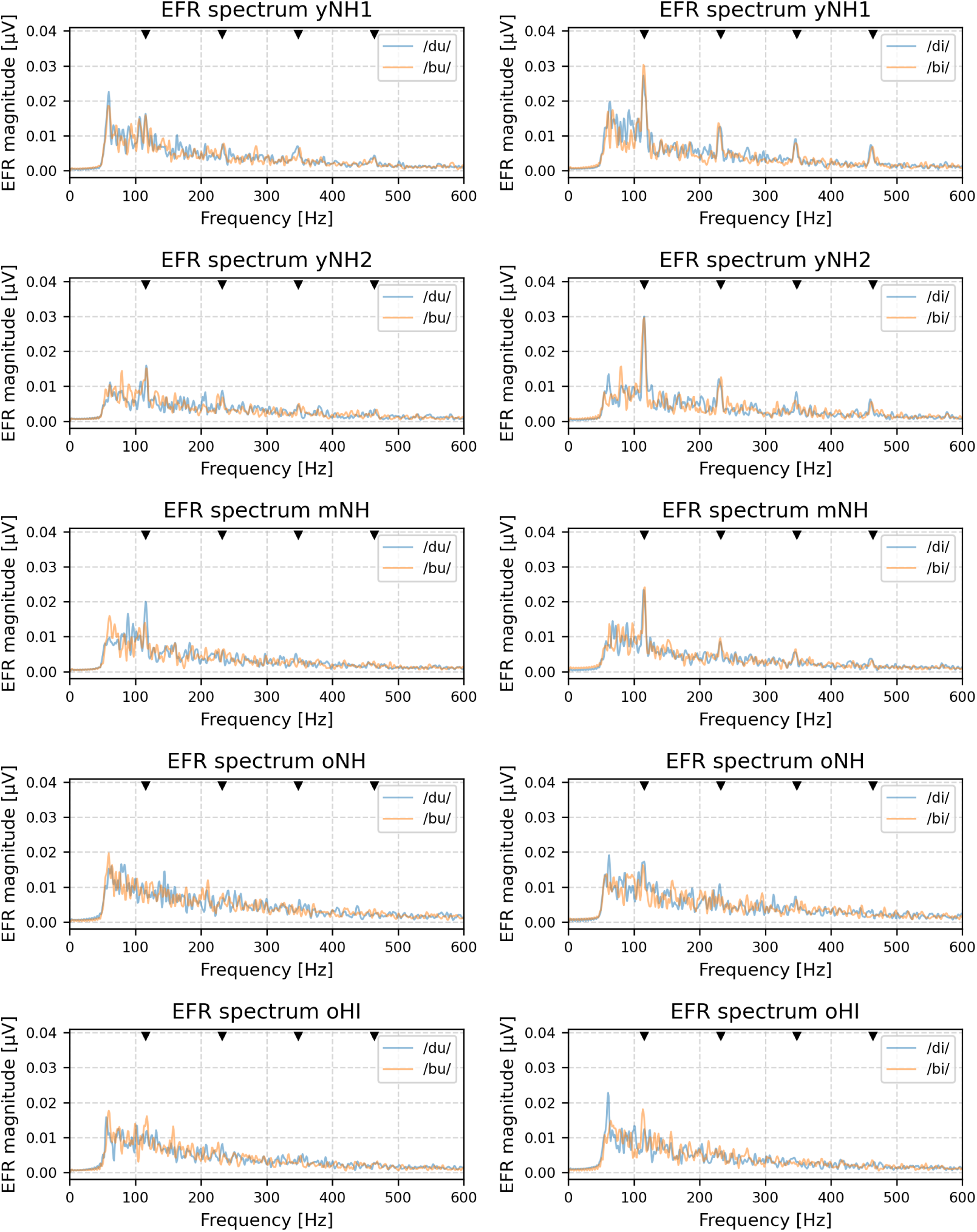
EFR frequency spectra for syllable contrasts for each group. The harmonics of *F*_0_ are marked by black triangle symbols.

**Figure A3:**
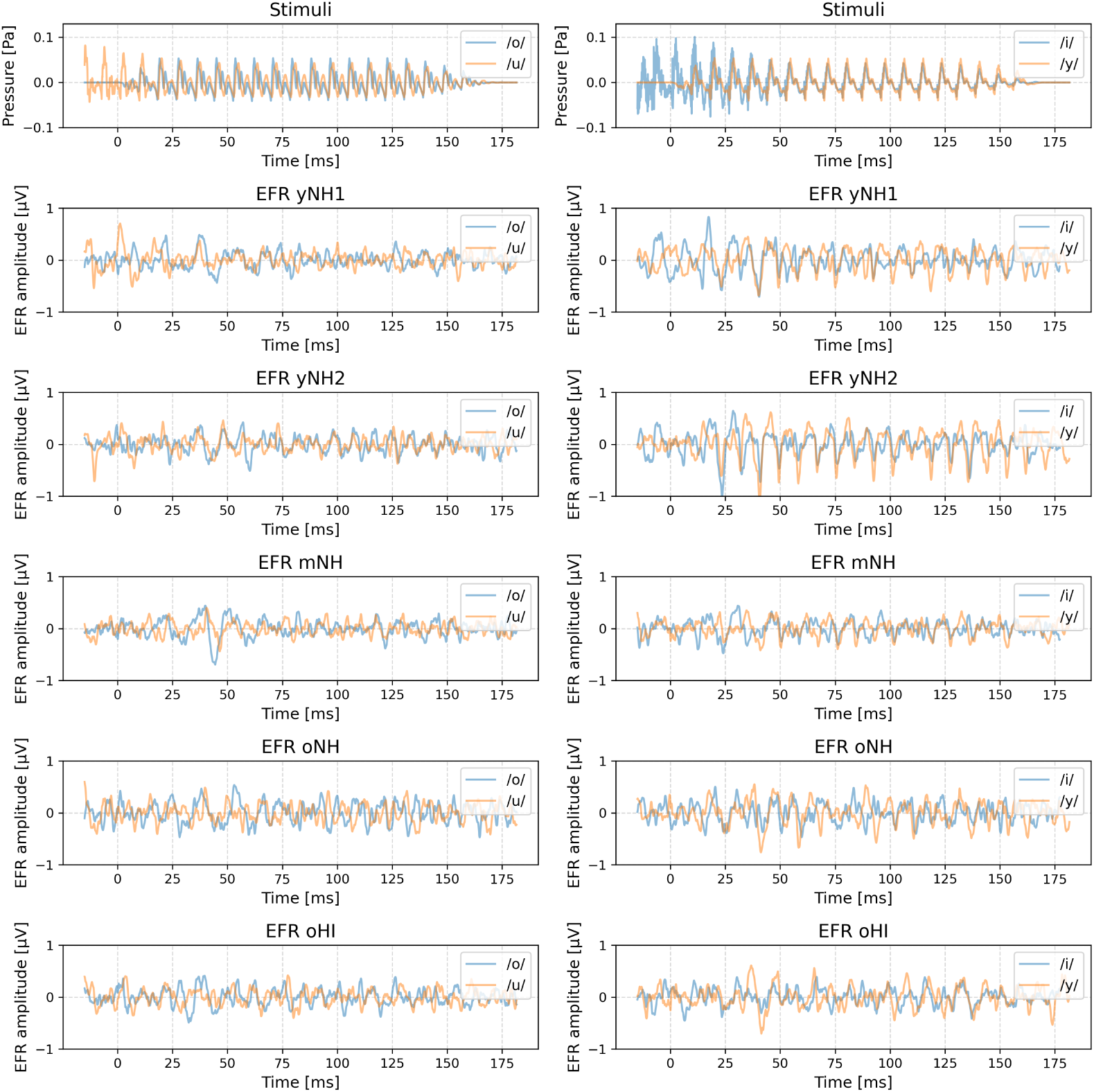
Time-domain waveform of the EFR for vowel contrasts for each group.

**Figure A4:**
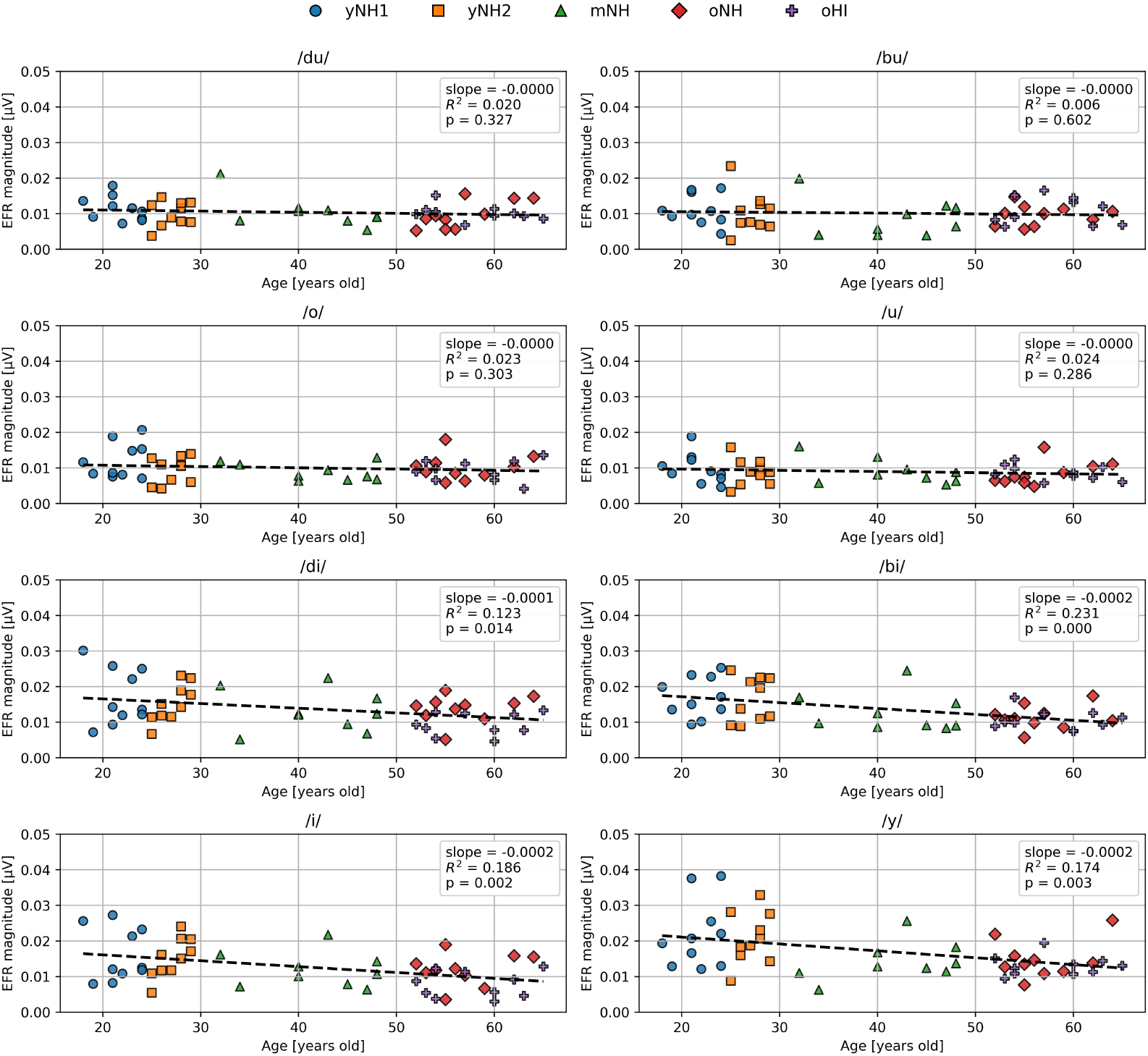
Scatter plots of EFR magnitude against age for each stimulus. The slope (*β*) and coefficient of determination (*R*^2^) of the linear regression were calculated across all groups, with associated p-values.

## Footnotes

1 There is an intense debate on the exact frequency of the PLL in humans (Verschooten et al., 2019). Depending on psychoacoustic tasks and measurement approaches, authors have estimated the PLL to be as low as 1.4 kHz and as high as 10 kHz. Our rather conservative choice of 1.5 kHz is based on two considerations: (i) our task involves neither pitch nor binaural perception, which are considered the most sensitive to TFS, and (ii) in the context of AEPs, phase-locked activity becomes increasingly impossible to distinguish from noise above 1 - 1.2 kHz (Skoe and Kraus, 2010). Despite the wide range of estimates reported by Verschooten et al. (2019), there is at least consensus over the fact that TFS availability decreases with frequency. Consequently, there is consensus on the fact that TFS should be more available below 1.5 kHz than above. However, it is worth keeping in mind that our interpretations are based on this conservative estimate and would have to be re-evaluated if new evidence proved this choice to be unrealistically low.

